# Network modeling of dynamic brain interactions predicts emergence of neural information that supports human cognitive behavior

**DOI:** 10.1101/2021.01.26.428276

**Authors:** Ravi D. Mill, Julia L. Hamilton, Emily C. Winfield, Nicole Lalta, Richard H. Chen, Michael W. Cole

## Abstract

How cognitive task behavior is generated by brain network interactions is a central question in neuroscience. Answering this question calls for the development of novel analysis tools that can firstly capture neural signatures of task information with high spatial and temporal precision (the “where and when”), and then allow for empirical testing of alternative network models of brain function that link information to behavior (the “how”). We outline a novel network modeling approach suited to this purpose that is applied to non-invasive functional neuroimaging data in humans. We first dynamically decoded the spatiotemporal signatures of task information in the human brain by combining MRI-individualized source electroencephalography with multivariate pattern analysis. A newly developed network modeling approach - dynamic activity flow modeling - then simulated the flow of task-evoked activity over more causally interpretable (relative to standard functional connectivity approaches) resting-state functional connections (dynamic, lagged, direct and directional). We demonstrate the utility of this modeling approach by applying it to elucidate network processes underlying sensory-motor information flow in the brain, revealing accurate predictions of empirical response information dynamics underlying behavior. Extending the model towards simulating network lesions suggested a role for the cognitive control networks (CCNs) as primary drivers of response information flow, transitioning from early dorsal attention network-dominated sensory-to-response transformation to later collaborative CCN engagement during response selection. These results demonstrate the utility of the dynamic activity flow modeling approach in identifying the generative network processes underlying neurocognitive phenomena.

## Introduction

Clarifying the spatial and temporal signatures underlying cognitive task information is critical to understanding the brain. Towards this aim, multivariate pattern analysis (MVPA) approaches have enabled decoding of different types of task information (e.g. sensory information in presented stimuli or response information underlying behavior) from evoked neural activation patterns (Haynes, 2015; Kriegeskorte and Kievit, 2013; Kriegeskorte et al., 2008). Prior research has shown these multivariate approaches to be more sensitive in relating neural activation measures to cognitive and behavioral variables of functional interest than earlier univariate approaches (e.g. single-cell spike rates, event-related potentials (ERPs) and functional MRI (fMRI) general linear model activations; Grootswagers et al., 2017; Jimura and Poldrack, 2012; Kriegeskorte et al., 2006; Saxena and Cunningham, 2019). These empirical observations underpin a shift in theoretical focus from localization of function to mapping distributed functionality at the neural population level (Eichenbaum, 2018).

Decoding methods applied to animal electrophysiological data have revealed that task-related information is highly distributed throughout cortex, such that many forms of task information can be decoded from multiple recorded brain regions (Bernardi et al., 2020; Kauvar et al., 2020; Raposo et al., 2014; Rigotti et al., 2013; Siegel et al., 2015). However, a degree of functional specialization has been demonstrated by scrutiny of temporal decoding characteristics, such as sensory information onsetting earliest in visual regions (Hernández et al., 2010; Siegel et al., 2015). Functional neuroimaging studies have interrogated the neural basis of task information non-invasively in humans, with fMRI highlighting a central role for large-scale spatial interactions between sensory/motor content networks and higher-order cognitive control networks (CCNs, i.e., dorsal attention, frontoparietal and cingulo-opercular networks; Cole et al., 2016a; Ito et al., 2017; Zhang et al., 2013). Recent applications of MVPA decoding in scalp/sensor-level electroencephalography/magnetoencephalography (EEG/MEG) data have hinted at complex information dynamics, involving short-timescale transitions between distinct whole-brain representations as sensory information is translated into a behavioral response (Gwilliams and King, 2020; Hubbard et al., 2019; King and Dehaene, 2014).

Critically, all of the aforementioned recording methods have their respective limitations: invasive electrophysiology has high spatial resolution, high temporal resolution but only partial spatial coverage; fMRI has high spatial resolution, low temporal resolution and full spatial coverage; sensor-level EEG/MEG has low spatial resolution, high temporal resolution and full spatial coverage. These methodological limitations have impeded a *comprehensive* spatiotemporal description of *both* spatially “where” and temporally “when” task information is decodable from brain activity. Moreover, the spatiotemporal coding schemes by which CCNs impact on task information and resulting behavior remain unresolved, as this would require analytic approaches that optimally balance spatial resolution, temporal resolution and whole-brain coverage.

Consequent to this lack of descriptive insight is an even greater dearth of understanding into the network computations (the “how”) underpinning task information. This deeper insight relies on the formalization of candidate models of how representations are computed (and transferred) across the brain, which are lacking in MVPA decoding approaches per se (de-Wit et al., 2016). To this end, it is likely that connectivity between neural entities plays a formative role in the computations that give rise to decodable information at specific spatial locations and temporal epochs (Ito et al., 2019). This follows from classical demonstrations of connectivity at the cellular level weighting the flow of action potentials via structural connectivity and Hebbian synaptic strength processes (Hebb, 1949; Levy and Steward, 1983). Such small-scale connectivity is theorized to coordinate the formation of “cell assemblies” representing task information at the population level (Caporale and Dan, 2008; Eichenbaum, 2018). A role for human neuroimaging-assessed functional connectivity (FC) as a large-scale aggregate of synaptic strengths has been hypothesized (Petersen and Sporns, 2015; Wig et al., 2011), and supported by evidence of long-term learning- and use-driven changes to FC (Lewis et al., 2009; Newbold et al., 2020a). Whilst this supports a role for repeated task coactivation in molding connectivity, a reciprocal influence is also anticipated by Hebbian theory: the intrinsic network architecture estimated via resting-state functional connectivity (restFC) should determine the likelihood of activity propagating between brain regions. Supporting evidence comes from recent “activity flow” models linking the emergence of task activations to communication pathways indexed by restFC. The observed accuracy in predicting empirical brain activations across a variety of cognitive tasks (Cole et al., 2016b; Ito et al., 2017) and in predicting dysfunctional activations and behavior in Alzheimer’s disease (Mill et al., 2020) substantiates the relevance of restFC in capturing the intrinsic network architecture that generates cognitively relevant phenomena (Mill et al., 2017).

However, these previous network models focused exclusively on predicting the spatial signatures of task activations, and overlooked accompanying dynamics. This is in part due to their sole application in fMRI which has well-known temporal limitations. We therefore developed a new dynamic version of activity flow modeling suited to higher temporal resolution EEG data (see Figure 1 for a task schematic and Figure 2 for our source modeling pipeline), so as to elucidate the dynamic neural computations underlying cognition. Importantly, we could not simply apply previous activity flow models – using contemporaneous brain activity in one region to predict contemporaneous activity in an independent brain region – to EEG data. This was due to well-known instantaneous field spread artifacts in EEG (and MEG) data (Schoffelen and Gross, 2009; Stinstra and Peters, 1998), which could result in analytic circularity if a to-be-predicted region leaked some of its signal into other regions used as predictors. A substantial innovation was necessary to overcome this limitation: updating activity flow modeling to use past activity to predict future activity. Given that it is impossible for brain activity to propagate back in time (a fundamental principle of causality and the direction of time), this considerably reduced the possibility of analytic circularity. As described in the Methods section, we implemented further rigorous steps during preprocessing (use of causal temporal filters), source reconstruction (use of dense-array EEG, individual structural MRIs and beamformer source modeling) and dynamic activity flow modeling (regressing out to-be-predicted region timeseries from all predictors prior to running all modeling analyses) to conclusively eliminate the risk of field spread and associated circularity.

**Figure 1.**
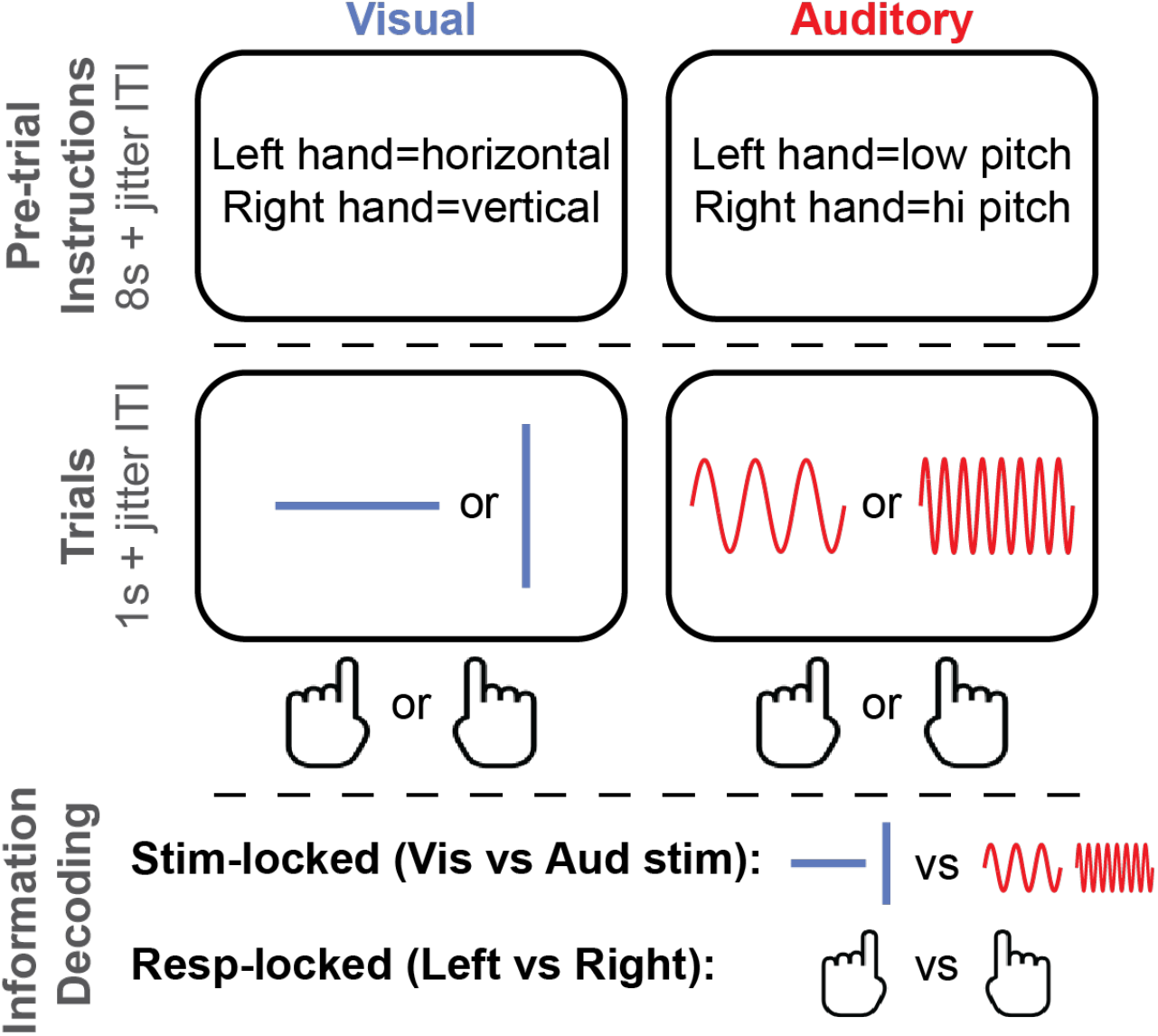
Task design. Depicted is one trial from each sensory block condition (10 trials per block, 12 blocks per subject session), and the types of decodable task information.

**Figure 2.**
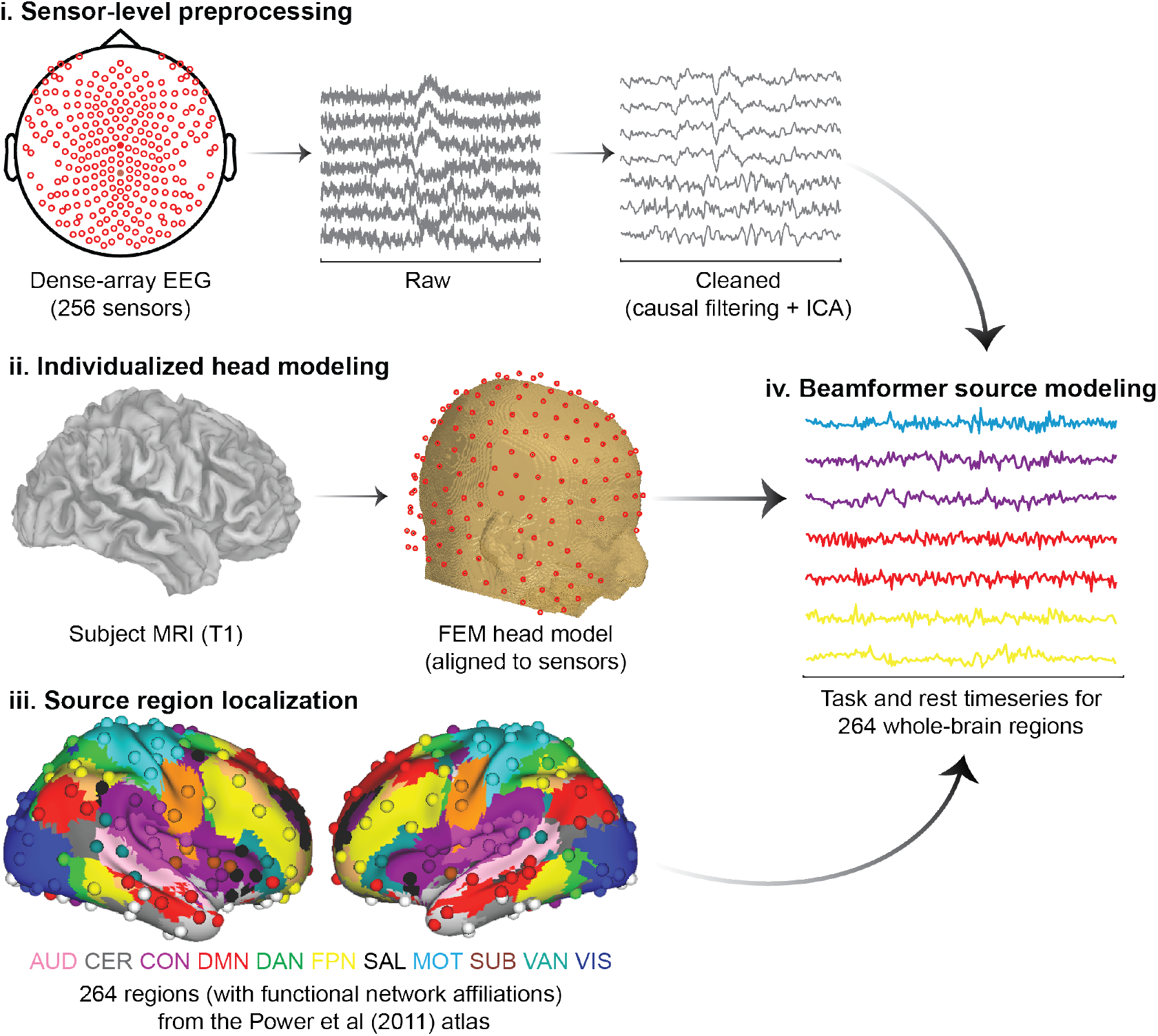
EEG preprocessing and source modeling pipeline. Applying this pipeline separately to task and resting-state EEG data reconstructed their respective activation timeseries. Colored text in panel iii) provides functional network affiliations for correspondingly colored regions localized from the Power functional atlas: AUD=Auditory; CER=Cerebellar; CON=Cingulo-opercular; DMN=Default mode; DAN=Dorsal attention; FPN=Frontoparietal; SAL=Salience; MOT=Motor; SUB=Subcortical; VAN=Ventral attention; VIS=Visual. See Methods for full details.

An added benefit of our novel network modeling approach was that the empirical restFC weights parameterizing the models were derived from multivariate autoregression (MVAR) applied in the resting-state (see Figure 3A). This captured dynamic, lagged, direct and directional connectivity between regions, rendering the models more causally valid than alternative approaches to estimating restFC (e.g. Pearson correlation, see Methods for how these features were derived from the causal inference/connectivity literatures). These innovations were necessary to achieve the desired prediction outcome: moment-to-moment fluctuations in future task information. This aligns activity flow modeling with the fully dynamic simulations of task information output by certain artificial neural network models (ANNs; Buonomano and Merzenich, 1995; Eliasmith et al., 2012; Yamins et al., 2014), with the added benefits of empirical estimation of connectivity weights (versus random initiation and optimization in ANNs; Sinz et al., 2019) and direct assessment of whether the engineered representations overlap veridically with those in the brain (Kriegeskorte and Douglas, 2018).

**Figure 3.**
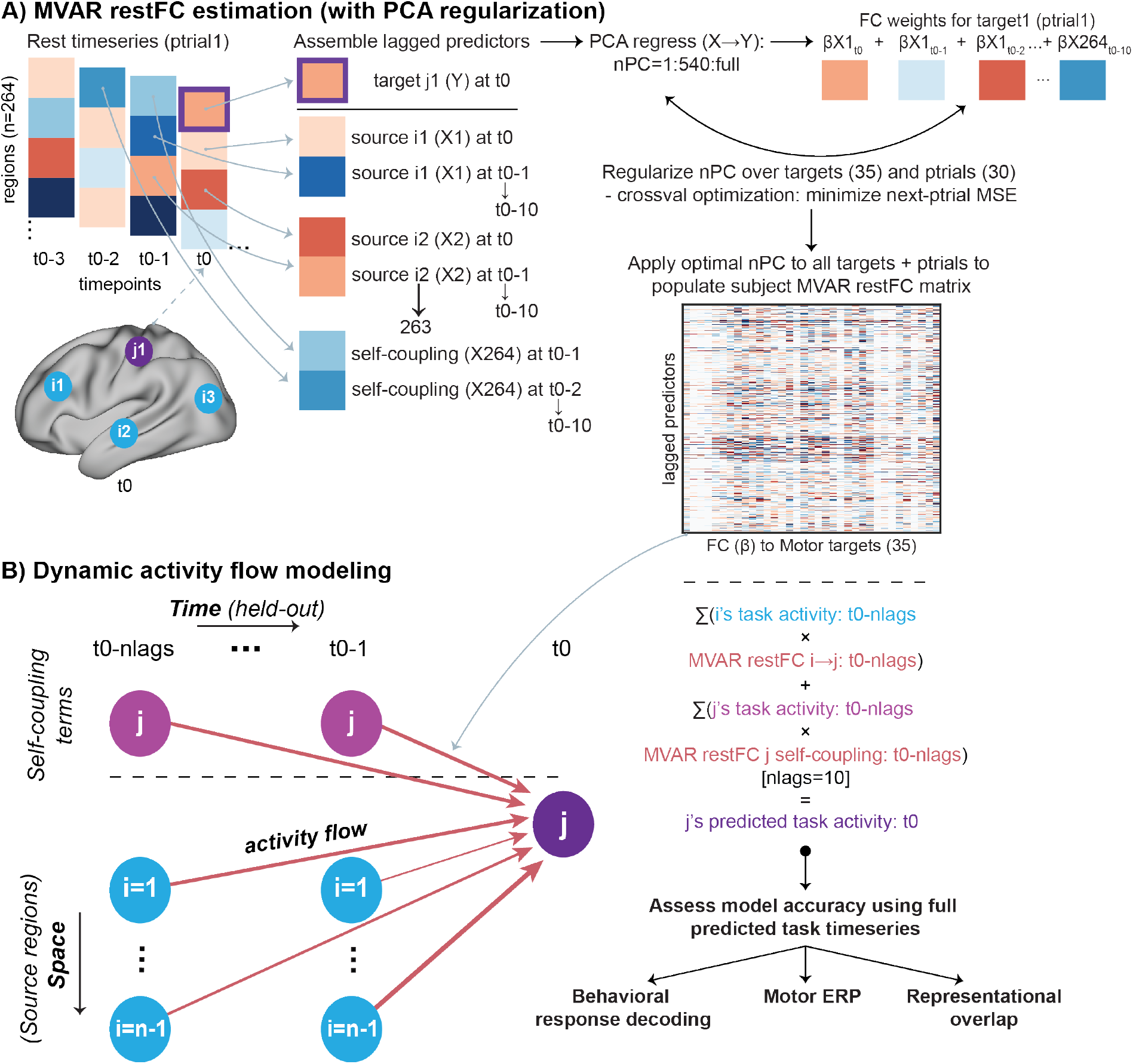
Approach to restFC estimation (via MVAR) and dynamic activity flow modeling. **A)** Estimation of MVAR restFC. Note that for demonstration purposes the schematic uses just 4 regions (1 Motor network target j1, and 3 predictor sources i1-3), whereas the full procedure iterated over 35 Motor targets and included all other 263 regions as sources. Lagged FC weights (β) from source regions and self-coupling terms to each Motor network region were calculated after regularizing the number of principal components (nPCs). This was achieved via crossvalidated minimization of the mean squared error (MSE) of the MVAR-predicted rest timeseries for held-out pseudotrials (ptrials). **B)** The lagged MVAR restFC weights were then combined with the lagged task activation timeseries to predict future Motor task activations via dynamic activity flow modeling. Iterating over all to-be-predicted Motor targets (35), trials (∼30) and trial timepoints (-0.45 to 0.45s around response commission) populated the full predicted Motor task activation matrix. This predicted matrix was the basis of subsequent response information decoding (dynamic MVPA), motor ERP and representational overlap analyses that assessed model accuracy.

In the present report, we sought to demonstrate the utility of this novel dynamic activity flow modeling approach via example application in clarifying the flow of cognitive task information in a simple sensory-motor categorization task (Figure 1). We first identified different forms of task information (sensory and response information, see Figure 1) using a combination of anatomically individualized EEG source modeling and dynamic MVPA, which richly described large-scale spatial and temporal information signatures (Figure 2).

After describing these signatures, we simulated their emergence using dynamic activity flow modeling (Figure 3). We first tested whether the full model can successfully predict future response information dynamics of the brain, thereby evidencing the cognitive and behavioral relevance of restFC as a large-scale analogue of synaptic strength processes governing neural information flow. Our ability to successfully model response information using connectivity weights that were entirely held-out from the task would also provide support for restFC in capturing the brain’s intrinsic (i.e. state-general) functional network architecture. Secondly, we extended the modeling framework to construct alternative network models (derived via simulated lesions to particular networks) to test the hypothesis that information flowing from the CCNs specifically (i.e. over and above the other functional networks) to the Motor network is central to producing behavior. We also sought to clarify the dynamic network computations used to fulfill this goal, so as to preliminarily highlight the functional utility of dynamic activity flow modeling in linking large-scale networks to more refined neurocognitive roles.

## Results

### Behavioral task performance

All subjects were able to perform the task, with accuracy significantly above chance for both the visual (mean=92.5%, Wilcoxon z=4.94, p<.001) and auditory (mean=91.5%, z=4.94, p<.001) conditions. Accuracy did not significantly differ between these two conditions (paired wilcoxon z=0.27, p=.789). The average reaction time for the visual and auditory conditions was 464.6s and 527.9s respectively, with RT significantly slower in the auditory condition (z=4.66, p<.001). This RT difference motivated a primary focus on response-locked rather than stimulus-locked trial data in subsequent analyses.

### Source modeling improves detection of cognitive task information

We compared peak information decodability for our approach that combined individualized source modeling and dynamic MVPA, with the more commonly used sensor-level dynamic MVPA approach. Timepoint-by-timepoint behavioral response information (correct left-vs right-handed responding) was decoded separately using all source regions (SourceAll) and all scalp sensors (SensorAll) as features. Visual inspection of the group response information timecourses (Figure 4) revealed multiple significantly decodable timepoints for both feature sets, with onset prior to commission of the response. This morphology is consistent with previous response decoding of EEG (Gwilliams and King, 2020) and invasive electrophysiological (Siegel et al., 2015) data.

**Figure 4.**
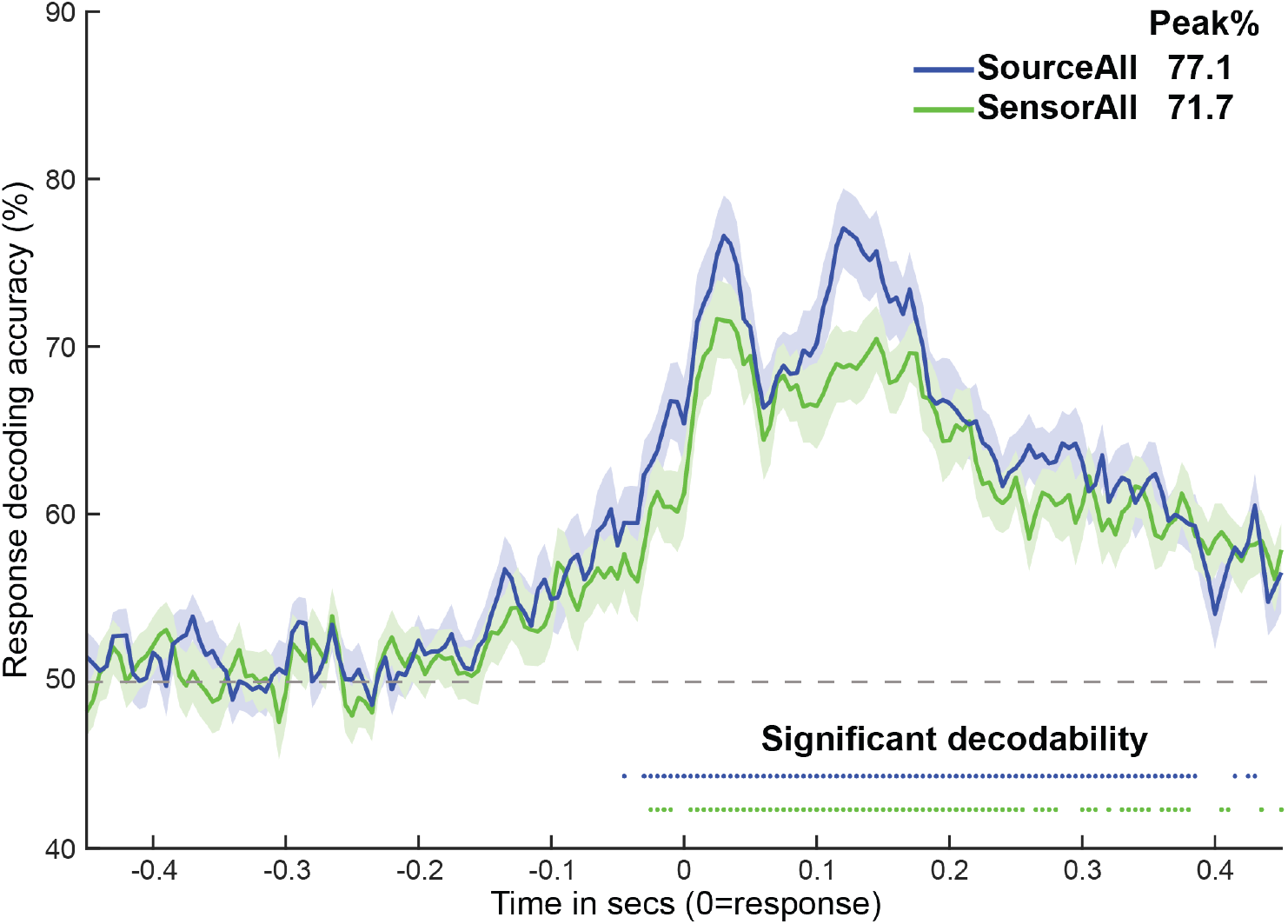
Detection of behavioral response information is improved for source versus sensor feature sets. Group-averaged response decoding timecourses for the SourceAll and SensorAll sets, with shaded patches reflecting the standard error of the mean across timepoints. Colored dots represent timepoints with significantly decodable information, as assessed by Wilcoxon signrank tests against 50% classification accuracy (p<.05, Bonferroni-corrected). The legend in the top right provides the peak decoding accuracy for each timecourse. Subject-level data underlying this figure are accessible via the public data repository (https://osf.io/mw4k3/, subdirectory: Results_figures_data/Figure4/).

Critically, peak response information decodability was visibly higher for the source-modeled approach (Figure 4). This was formalized statistically by extracting peaks from each subject’s SensorAll and SourceAll decoding timecourses, and contrasting them via paired Wilcoxon tests. The SourceAll decoding peak was significantly higher than the SensorAll peak across variation in how time-to-peaks were defined: i) from the SensorAll group time-to-peak (0.025s; 3.8% peak change; z=2.47, p=.014), ii) from the SourceAll group time-to-peak (0.120s; 8.4% peak change; z=3.57, p<.001), iii) from the peak of each individual subject’s decoding timecourse (unbiased by the group results; 4.8% peak change; z=3.59, p<.001). Peak decodability of sensory information (visual vs auditory stimulus condition) was also numerically higher for SourceAll compared to SensorAll features (see SI, Figure S1A). Hence, combining source modeling with dynamic MVPA significantly improved detection of response information. This might have arisen from improvements in signal-to-noise introduced by beamformer source modeling (Brookes et al., 2008), and opposes prior suggestions that beamforming leads to overarching cancellation of the underlying neural activations (Hui et al., 2010; Van Veen et al., 1997). These results extended our critical requirement for subsequent analyses - that spatially distinct regions were localized accurately - by demonstrating that such source localization can actually improve overall decodability.

### Network decoding reveals prominent roles of Motor and Cognitive Control Networks in representing behavioral response information

We then applied our source-modeled dynamic MVPA approach to decode spatial and temporal signatures of response information from each of the 11 major functional networks (Power et al., 2011), treating within-network regions as features (see Methods). To clarify, we focused on the network rather than region level as a principled decision following prior demonstrations of the precision afforded by EEG source modeling (∼3-10 mm cortical localization error; Klamer et al., 2015; Lascano et al., 2014; Seeber et al., 2019). The lower end of this range questions an alternative “searchlight” approach previously applied to fMRI data, which decodes information using within-region voxels/vertices (typically of <∼3mm resolution) as features. We hence focused on the network level, which aligned with our a priori interest in making functional inferences at this level. Despite this spatial scale being somewhat coarse (compared to invasive animal methods), it is worthwhile reiterating that this permits a degree of spatial insight that sensor-level EEG/MEG analyses are singularly incapable of.

The network decoding results are depicted in Figure 5. Consistent with the morphology of the SourceAll response decoding timecourse (Figure 4), we again observed two decoding peaks across networks. In the Supplementary Information, we link this two-peak morphology to sequential motor preparation and motor execution/feedback processes, both via stimulus-locked response decoding (Figure S3A and Figure S3B) and via a temporal generalization analysis that revealed distinct multivariate codes for each peak (Figure S3C).

**Figure 5.**
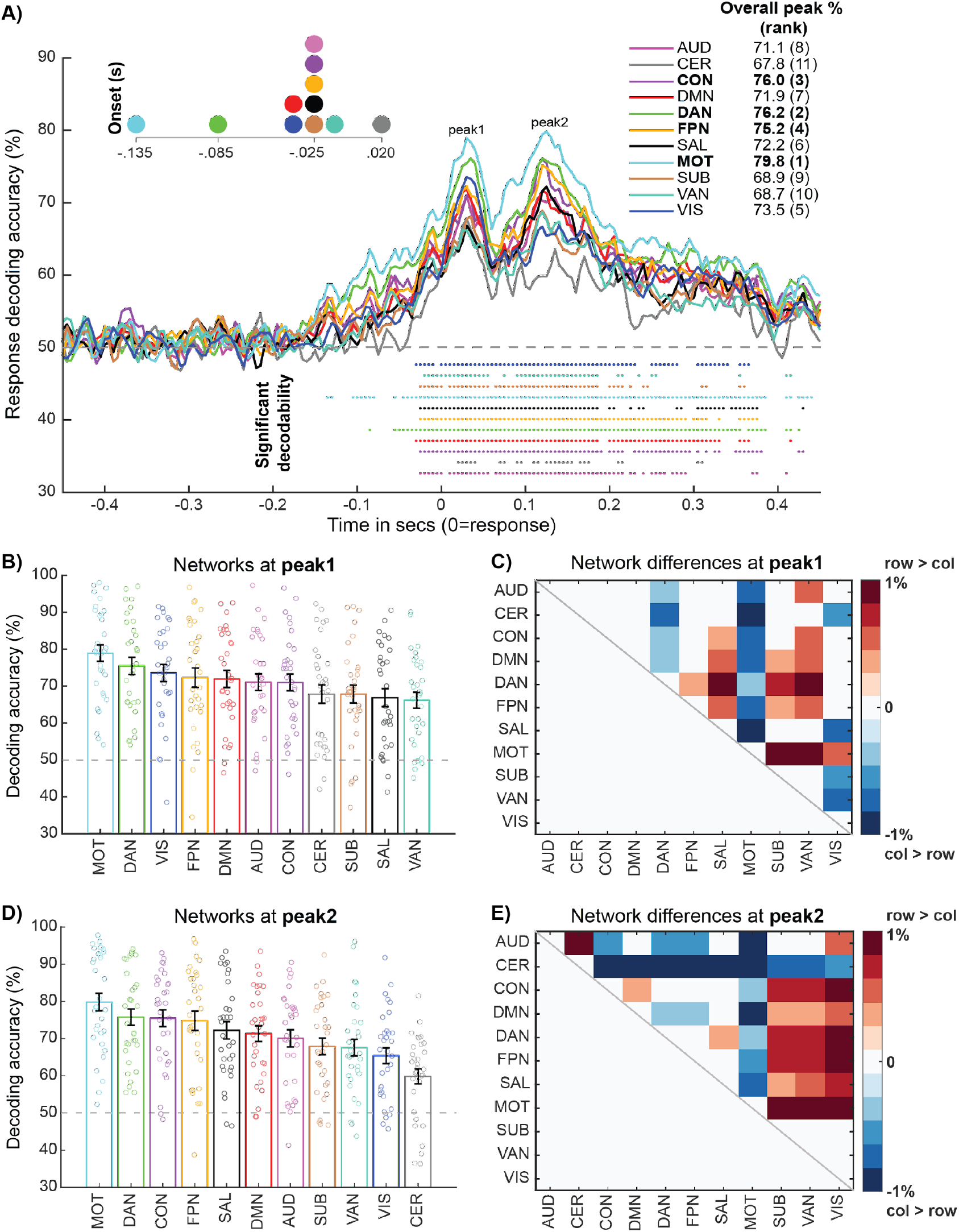
Network decoding of behavioral response information reveals prominent roles for Motor and Cognitive Control networks. **A)** Group decoding timecourses color-coded by network affiliation. Colored dots represent significantly decodable timepoints for each network (p<.05 via Wilcoxon signrank against 50% chance, Bonferroni-corrected) and the legend in the top right provides peak decoding accuracies for each network. A magnified plot of the onset of the first significant timepoint for each network is provided in the top left. **B)** Response decoding accuracy ranked across networks at peak 1 (0.03s). Each bar represents the mean and standard error for each network, with individual subject data points also overlaid. **C)** Matrix capturing the significance of cross-network differences in decoding accuracy at peak 1. Plotted is the pairwise difference in mean decoding accuracy, thresholded via paired Wilcoxon tests (p<.05, FDR-corrected). Positive values denote significantly higher decoding accuracy for the row network > the column network, and vice versa for the negative values. **D)** and **E)** follow the same conventions as C) and D) respectively, albeit focusing on peak 2 (0.125s). Subject-level data underlying this figure are accessible via the public data repository (https://osf.io/mw4k3/, subdirectory: Results_figures_data/Figure5/).

Response information was broadly decodable across networks, consistent with prior findings from primate electrophysiology (Bernardi et al., 2020; Siegel et al., 2015) and rodent optical imaging (Kauvar et al., 2020). However, despite the visible similarity of the network timecourses, examination of timecourse features revealed content-appropriate specialization: response information onset earliest and peaked strongest in the Motor network. Similar content-appropriate specialization was observed when decoding stimulus categories within each sensory modality, with visual stimulus information (horizontal versus vertical lines) peaking in the Visual network and auditory stimulus information (low versus high pitch sounds) peaking in the Auditory network (see SI, Figure S2).

Prominent response decodability was also observed in the CCNs, all three of which yielded the highest response information peaks after the Motor network. We statistically formalized these between-network effects by contrasting decoding timecourse peaks extracted from individual subjects (see Methods). We extracted subject-level peaks separately for peak 1 (motor preparation, Figure 5B) and peak 2 (motor execution/feedback, Figure 5D), from the respective cross-network group peak timepoints within these windows (0.03s and 0.125s). Motor network activity was decoded significantly higher relative to other networks across both peak 1 (Figure 5C) and peak 2 (Figure 5E). Interestingly, whereas Visual network activity was prominently decoded during peak 1 (see network ranking in Figure 5B), it was much less involved in peak 2 (Figure 5D). One possibility is that this earlier period reflects motor preparation operations that translate sensory into response information, whereas the later peak 2 period may reflect purer motor representations underlying response execution and feedback, consistent with prior animal (Elsayed et al., 2016; Raposo et al., 2014) and human motor ERP (Wascher and Wauschkuhn, 1996) research.

The transition from peak 1 to peak 2 also charted an interesting differentiation in the engagement of the CCNs. The DAN was more prominently involved in the motor preparation peak 1, and differed significantly from both the CON and FPN (Figure 5B). This profile of preferential DAN engagement was also observed when sensory information was more directly decoded from the stimulus-locked data (see SI, Figure S1B), suggesting that similar sensory-related computations might be occurring at peak 1 in the response timecourse. However, the later peak 2 elicited a different network profile: information was higher for all three CCNs relative to the other networks but was not reliably differentiated between them (even at an uncorrected threshold of p<.05, Figure 5E).

Overall, the network decoding results highlight the utility of decoding information from neuroimaging data that is well-resolved in both the spatial and temporal domains, as was uniquely enabled by our combination of source EEG and dynamic MVPA. Beyond recovering the content-appropriate specialization of the Motor network for response information, this approach led to a segregation of dynamic CCN profiles: from preferential engagement of the DAN during translation of sensory to response information at peak 1, to more collaborative engagement of all three CCNs to generate purer response representations at peak 2. In the Supplementary Information (see SI, Figure S4), we demonstrate that the pattern of network decoding results is highly similar when using an alternative approach that confines the decoding to more “unique” signals within each network.

### Dynamic activity flow modeling predicts future response information dynamics

The preceding network decoding results described which spatial networks prominently represent response information (“where”) and their accompanying temporal profiles (“when”). We next sought insight into “how” these representations emerge computationally from dynamic network interactions in the brain. We treated this simplified sensory-motor task scenario as a testbed for our novel network modeling approach (dynamic activity flow), and used it to predict future response information dynamics from past task activations flowing over communication pathways parameterized by restFC. Based on the content-appropriate specialization of the Motor network demonstrated above (Figure 5), we considered regions within this network as to-be-predicted “targets” for the emergence of neural representations driving overt behavior.

Functional connectivity to these regions from the rest of the brain was estimated via multivariate autoregression (MVAR) in a separate resting-state EEG session. This MVAR approach was constrained via principles adapted from the causal inference (Pearl and Mackenzie, 2018) and causal connectivity literatures (Mumford and Ramsey, 2014; Ramsey et al., 2010; see Methods for details) to output dynamic, lagged, direct and directional estimates of restFC (see Figure 3A). The restFC regression weights for each subject were appropriately regularized via optimized PCA regression^1^.

These lagged restFC weights were then combined with lagged task activations to generate future model-predicted activation timeseries for all Motor network regions across the entirety of the task (Figure 3B). The Supplementary Information depicts the correlation-based group restFC matrix and presents some descriptive analyses verifying recovery of the canonical functional network architecture (as a sanity check), as well as the inter-subject reliability of the MVAR FC weights (see SI, Figure S5).

We first assessed the accuracy of the dynamic activity flow model in capturing the emergence of behavioral response information in this simple categorization task. The predicted Motor region activation timeseries underwent the same dynamic MVPA procedure applied previously to the actual data. This yielded a predicted Motor network response information timecourse, which is depicted in Figure 6A along with the actual Motor timecourse. The plot demonstrates that overt behavioral information was decodable at multiple timepoints from the predicted data which, to reiterate, was generated exclusively from FC weights derived from a held-out rest session, and held-out (past) temporal task information. The predicted decoding peak was in fact significantly higher in magnitude (82.3%) than the actual peak (79.8%), as formalized by a paired Wilcoxon contrast of the subject-level peaks (extracted from the actual group time-to-peak at 0.125s): z=2.36, p=.018. We advocate caution in inferring too much from this greater decodability in the predicted data (which, as a 2.5% peak difference, is numerically small), and in subsequent sections (accompanying Figure 6B and 6C) detail more intuitive findings of greater significance in the actual compared to the predicted data. These findings suggest that this singular instance of greater decodability in the predicted data likely arose from a trivial source, such as partial overfitting to noise.

**Figure 6.**
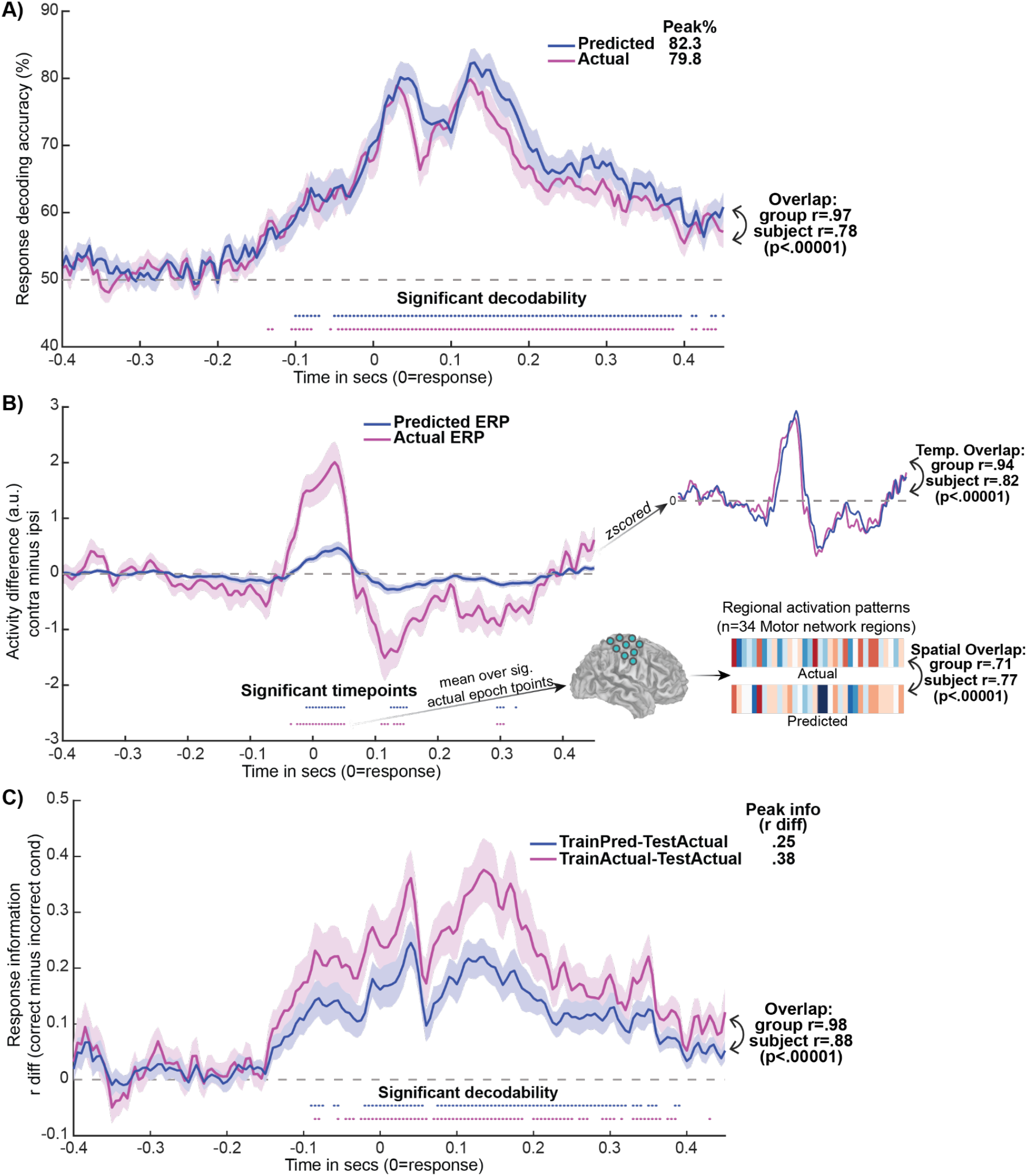
Dynamic activity flow modeling accurately and veridically predicts future behavioral response information. **A)** The model accurately predicted future response information dynamics in the Motor network. Predicted and actual group decoding timecourses are plotted, with overall decoding peaks (top right), and predicted-to-actual timecourse overlap (Pearson r) at group and subject levels. **B)** The motor ERP waveform (and spatial pattern) was also successfully predicted, demonstrating fidelity to the underlying activations.Group averaged ERP difference waves are plotted (contralateral minus ipsilateral; a.u.=arbitrary units). To aid visualization, the top right depicts the zscored group waveforms, along with the temporal predicted-to-actual overlap (at group and subject levels). The bottom right provides the spatial overlap between the predicted and actual Motor region activation vectors, extracted over the -0.035 to 0.050s epoch of significance in the actual data. C) Representational overlap analyses highlight veridicality of the model-predicted representations, given their ability to decode the actual data. The group information timecourse resulting from training on predicted representations and testing on the actual data (TrainPred-TestActual) is depicted, as well as the result of training and testing on the actual data (TrainActual-TestActual) for comparison. Response information was quantified as the difference in Pearson r values computed for the test activation pattern at each timepoint with correct minus incorrect response condition templates (see Methods). Overall information peaks (top right) and temporal overlap between the two timecourses (bottom right) are provided. For all panels, significance at each timepoint was assessed via Wilcoxon signrank tests (versus 50% chance for panel A, and versus 0 for B/C) with Bonferroni correction. Subject-level data underlying this figure are accessible via the public data repository (https://osf.io/mw4k3/, subdirectory: Results_figures_data/Figure6/).

Critically, comparison of the overlap between the predicted and actual decoding timecourses confirmed that the model faithfully captured the temporal morphology of the information timecourse. This held at the group-level (r=.97, p<.00001) and also at the subject-level (r=.78, p<.00001); further emphasizing the accurate individualized predictions obtained. The subject-level coefficient of determination (R^2^, capturing scaled prediction accuracy, see Methods) was also reliably greater than 0 (mean R^2^=0.42, p<.00001), indicating that dynamic activity flow modeling outperformed a null model built from the mean. Similarly high model prediction accuracy was achieved when using more common non-causal filtering during preprocessing (see SI, Figure S6). In the Supplement we also highlight the utility of our rigorous regression approach to controlling for field spread artifacts, which was applied in all modeling analyses reported here to effectively control for potential leakage/circularity (see SI, Figure S7).

Finally, we also compared the accuracy of our model’s predictions to a permuted null model in which predicted response information timecourses were generated (over 500 permutations) after scrambling the MVAR FC terms. Critically, the order of the autoregressive activation terms were preserved, to provide a targeted test of the informativeness of the FC terms specifically in accurately simulating response information. Comparing the proportion of permuted prediction accuracies greater than that observed in the unscrambled data confirmed that the FC terms were highly informative (observed r > permuted r, p=.002, at group and subject levels).

The predicted dynamic MVPA results hence attest to the fidelity of the dynamic activity flow model in capturing the network-weighted flow of response information in a highly temporally resolved and individualized fashion. This highlights the utility of the modeling approach in evidencing the cognitive relevance of restFC in generating response information dynamics underlying behavior.

### Dynamic activity flow modeling also predicts the raw activation timeseries (motor ERP)

We then probed the model’s accuracy in predicting activations in Motor network regions that were the basis of the dynamic MVPA analysis. This would demonstrate fidelity to the underlying neural activation timeseries, and challenge the possibility that the strong response information decoding in the predicted data emerged from factors artificially introduced by the model.

We targeted accurate prediction of an established motor event-related potential (ERP) termed the lateralized readiness potential or bereitschaftspotential (Cheyne et al., 2006; Deecke et al., 1976; Kutas and Donchin, 1980): greater activation at the time of response commission for Motor network regions that are contralateral to the response hand. The predicted and actual group motor ERP waveforms are provided in Figure 6B (as contralateral minus ipsilateral difference waves, see Methods). Multiple significant activation timepoints were observed in the actual and predicted timecourses, albeit these were anticipatedly greater in number for the actual data. Critically, high overlap in the temporal morphologies of the predicted and actual timecourses was observed at both the group and subject levels (r=.94 and r=.82 respectively, both p<.00001). We also probed the predicted-to-actual overlap in the spatial domain, by correlating the respective activation vectors for Motor network regions averaged across the epoch of significance in the actual data (-0.035 to 0.050s, Figure 6B). This revealed a high degree of spatial overlap between the predicted and actual spatial activation patterns, at the group (r=.71) and subject levels (r=.77, both p<.00001). This high spatial overlap held when the predicted and actual vectors concatenated timepoints across the entire trial, at both the group (r=.70) and subject levels (r=.61, both p<.00001). Subject-level R^2^ values were also reliably greater than 0 indicating superior model performance than the mean, across temporal (mean R^2^=0.25, p<.00001) and spatial (mean R^2^=0.16, p<.0001) overlap domains.

Overall, the motor ERP results illustrate that the dynamic activity flow model maintained high fidelity to the actual task activations; across both temporal and spatial domains, and even at the individual subject level.

### The model is directly faithful to the veridical representational geometry

Whilst the previous section reported highly accurate predictions of the underlying Motor network activations, we sought to more directly demonstrate that the strong response information decoding obtained via dynamic activity flow modeling was driven by multivariate representations that overlapped with how the brain veridically represents response information. This aligns with mounting emphasis on the need to optimize representational overlap between neural network models and the brain (Kriegeskorte and Douglas, 2018; Yamins and DiCarlo, 2016; Yang and Wang, 2020).

To address this, we investigated whether actual response information in the Motor network could be dynamically decoded from the model-predicted representations. The group information timecourses are presented in Figure 6C: 1) for trained representations in the predicted data applied to decode the actual data (“TrainPred-TestActual”), and 2) for trained representations in the actual data applied to decode the actual data (“TrainActual-TestActual”). The figure highlights multiple significantly decodable timepoints in the key TrainPred-TestActual timecourse, providing direct evidence that the dynamic activity flow model captured neurally valid multivariate representations. In keeping with the preceding ERP analyses, the number of significantly decodable timepoints (and the overall information peak) was anticipatedly higher in the TrainActual than the TrainPred model. Critically, the morphology of the resultant response information timecourse also temporally overlapped with that obtained for the TrainActual-TestActual model, at the group and subject level (r=.98 and r=.88 respectively, both p<.00001). Subject-level R^2^ was also reliably greater than 0 (mean R^2^=0.59, p<.00001). This suggests that the dynamics in representational geometry over the entire trial were well captured. Overall, this representational overlap analysis provides compelling evidence that the dynamic activity flow model preserves high fidelity to the veridical neural codes underlying response information.

### Model-simulated network lesioning suggests prominent influence of the CCNs in driving response information flow

We modified dynamic activity flow modeling to provide additional insight into the network computations that generate behavior. Separate models predicting response information in the Motor network were run that systematically lesioned all but one of the remaining functional networks, whilst also excluding the autoregressive/self-coupling terms (see Methods). Beyond this iterative restriction of source predictor terms (Figure 3B), the procedure was identical to that detailed in the dynamic MVPA analysis of the model-predicted data. This yielded 10 response decoding timecourses for each functional network model, capturing their putative influence on information flow to the Motor network. We then adopted a model comparison approach (similar to earlier network modeling approaches e.g. Penny et al., 2004) to identify which networks were the most prominent drivers of response information flow to the Motor network in this simple task, via rigorous statistical comparison of the information peaks across networks with multiple comparison correction.

The network-lesioned group decoding timecourses are presented in Figure 7A. Visual inspection revealed significant decodability for all network models, however this peaked highest for the CCN models (top right Figure 7A), with onset also preceding the other networks (top left Figure 7A). The timecourses once again followed a 2-peak morphology, with CCN-derived decodability visibly varying across the earlier motor preparation epoch and the later motor execution/feedback epoch (as in Figure 5).

**Figure 7.**
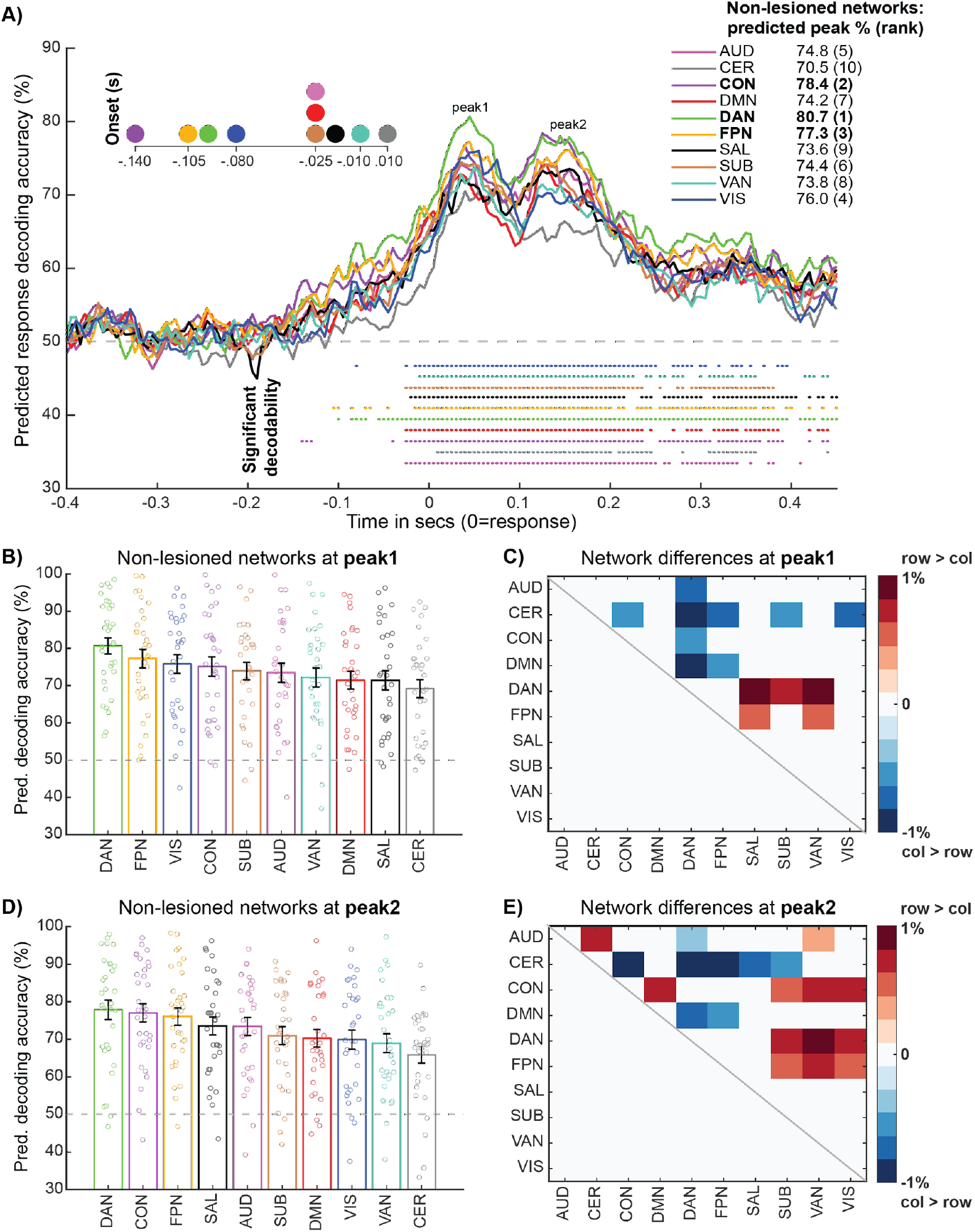
Network lesioning reveals contributions of individual functional networks in driving future behavior. Unlike the visually similar Figure 5 that provided descriptive insight, these analyses provided explanatory insight into which spatial networks likely drive response information representations in the Motor network, and their accompanying temporal signatures. For these analyses all networks were lesioned except for the indicated (non-lesioned) network. **A)** Group predicted decoding timecourses for each of the network-lesioned models, color-coded by affiliation as before. Magnified decoding onsets (top left), overall decoding peaks (top right) and significant decodability at each timepoint (assessed via bonferroni-corrected Wilcoxon tests, as before) are provided for each network model. **B)** Predicted response decoding accuracy ranked across network models at peak 1 (0.045s). Each bar represents the mean and standard error for each network, with overlaid subject data points. **C)** Matrix capturing cross-network differences in predicted decoding accuracy at peak 1. Plotted is the pairwise difference in mean decoding accuracy, thresholded via paired Wilcoxon (p<.05, FDR-corrected). Positive values denote significantly higher predicted decoding accuracy for the row network > the column network, and vice versa for the negative values. **D)** and **E)** follow the same conventions as C) and D) respectively, except now focusing on peak 2 (0.155s). Subject-level data underlying this figure are accessible via the public data repository (https://osf.io/mw4k3/, subdirectory: Results_figures_data/Figure7/).

However, these network lesioning analyses critically extended the descriptive results in Figure 5, highlighting that whilst the CCNs exerted a broadly strong predictive influence across the trial, this pattern of influence differed across the two peaks. These differing network profiles were statistically formalized by between-network contrasts (model comparisons) of subject-level predicted decoding peaks, which were extracted separately from identifiable group peak timepoints for peak 1 (0.045s) and peak 2 (0.155s). At the motor preparation peak 1 (Figure 7B), the DAN network model yielded response decodability that was significantly higher than the CON (p<.05 FDR-corrected, Figure 7C) and marginally higher than the FPN (p<.05 uncorrected). Considering the strong influence of the Visual network also observed at peak 1 (Figure 7B), the results again support more selective engagement of the DAN during this likely period of sensory-related processing (e.g. translating sensory into response information). At the later motor execution/feedback peak (Figure 7D), all three CCNs exerted the strongest generative influence on response decodability, and were not reliably differentiated amongst each other (Figure 7E), even at lenient p<.05 uncorrected thresholds. The pattern of network model differences again suggests more collaborative engagement across the CCNs, possibly reflecting the formation of purer response representations during this later period.

The network lesioning results therefore extended previous description of spatiotemporal signatures underlying response information to elucidate the generative influence of the CCNs on network information flow driving overt behavior. This prominent influence of the CCNs was also recovered when using subject-level prediction accuracy for each network model as an alternative metric to the predicted decoding peaks used above (see SI, Figure S8).

### Actual and model-predicted results hold with reduced numbers of EEG electrodes

We explored the effect of varying analysis parameters on the results. Firstly, we reduced the numbers of available EEG electrodes by confining the preprocessed causal filter data to standard 64- and 128-electrode montages, to test the generalizability of our source modeling and dynamic activity flow network modeling approaches across available EEG systems. To derive these montages, the spatial euclidean distance between template coordinates for each electrode in the 64-/128-electrode systems and electrode coordinates in the original 256-electrode system was computed.

The 256 system electrode with the minimum euclidean distance with a given electrode in the 64/128 system was uniquely assigned to it.

For the SourceAll vs SensorAll response decoding analysis, we recovered the original finding of higher peak information decodability for source data across both reduced electrode montages (see Figure 8A, which includes the results for the 256 system for reference). However, the group information peak in the source data increased with increasing numbers of electrodes (see panel in Figure 8A). This conclusion was quantified by extracting unbiased information timecourse peaks at the subject level (as before), and submitting them to a 3 (electrodes: 64, 128, 256) x 2 (data type: sensor, source) repeated measures ANOVA. This revealed a main effect of data type that confirmed the superiority of source compared to sensor information decoding across electrode numbers (subject source peak mean=89.0%, sensor peak mean=84.9%, F(1,31)=29.29, p<.001, n^2^G=0.079). The main effect of electrodes was non-significant (F(2,62)<1). We also observed a non-significant interaction, suggesting a trend towards increasing source versus sensor superiority with increasing number of electrodes (F(2,62)=2.90, p=.063, n^2^G=0.003). Planned paired Wilcoxon contrasts of source peaks across the three electrode types were non-significant, but suggested a numerical ‘jump’ in source decodability for 128 and 256 systems compared to the 64 system (source 128 vs source 64, z=1.65, p=.102; source 256 vs source 64, z=1.70, p=.092; source 256 vs source 128, z=0.80, p=427). Hence, the central finding of greater source compared to sensor decodability held with reduced numbers of electrodes. This complements the more fundamental benefit of source modeling in allowing for more valid spatial inferences (a prerequisite for our network decoding and dynamic activity flow modeling analyses), whereas sensor-level analyses are inherently spatially ambiguous. We also found a numerical (albeit in this case statistically unreliable) increase in the magnitude of this source advantage for systems with greater than 64 electrodes.

**Figure 8.**
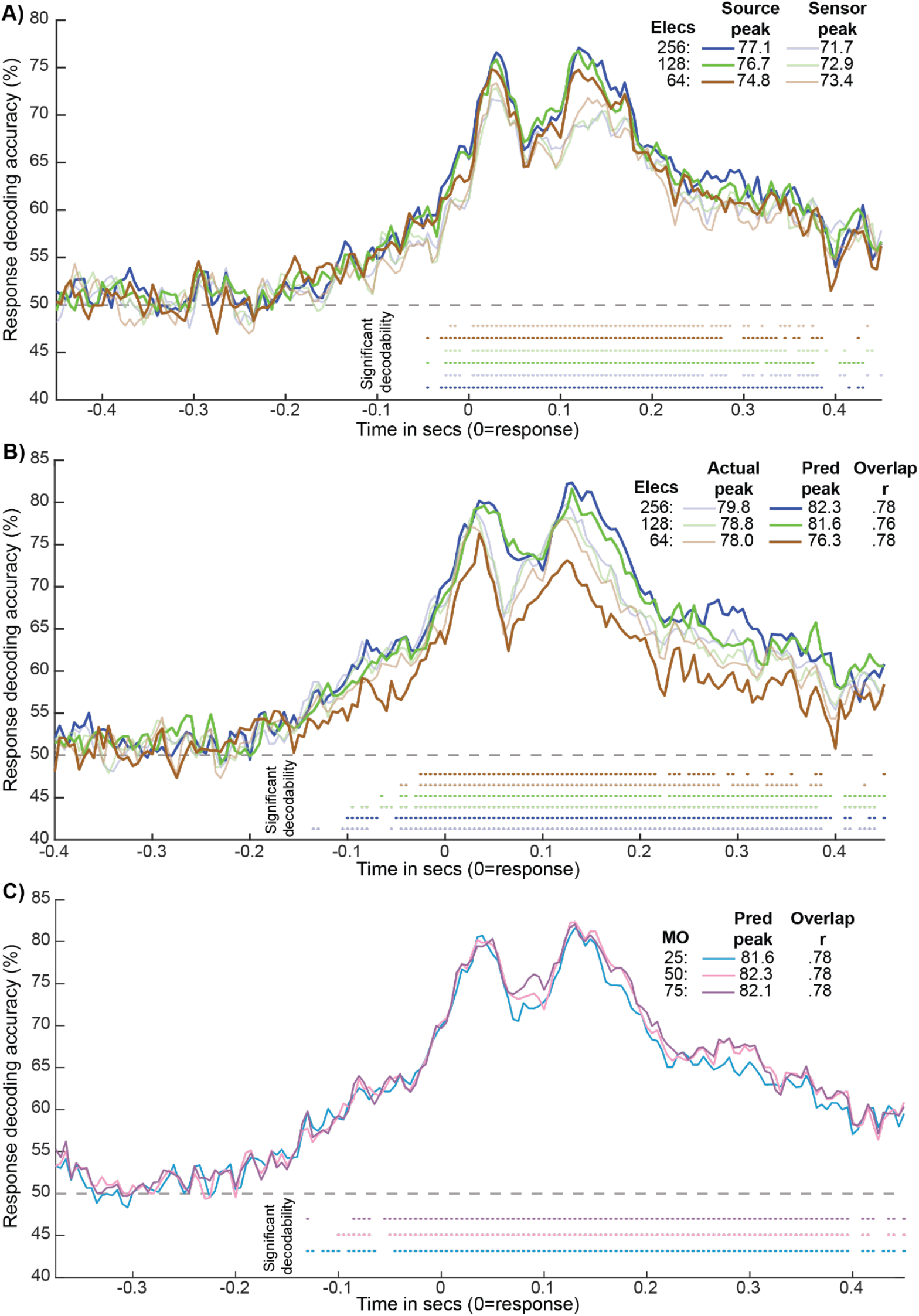
Control analyses varying number of electrodes and model order. **A)** Group response decoding timecourses for the SourceAll (dark lines) and SensorAll (softer lines) feature sets, plotted across variation in number of available electrodes (color-coded). **B)** Group response decoding timecourses for actual (softer lines) and model-predicted (darker lines) Motor network information, plotted across electrode systems. Note that subject-level predicted-to-actual overlap r values are also provided in the legend (all p<.00001). **C)** Group response decoding timecourses for model-predicted Motor network information across variation in model order (number of lagged predictors). Subject-level predicted-to-actual overlap r values are also provided in the legend (all p<.00001). For all panels, significantly decodable timepoints and group peak decoding accuracies are depicted per conventions in the earlier figures.

We also repeated the main dynamic activity flow modeling analysis for the reduced 64- and 128-electrode systems. Figure 8B plots actual and model-predicted Motor network response information timecourses across the 3 electrode systems. Again, we find that the main modeling results hold even when fewer electrodes are available: dynamic activity flow modeling reliably predicted response information (multiple significantly decoded timepoints in the predicted timecourses) and accurately captured the temporal morphology (significant subject-level predicted-to-actual overlap r values, see panel of Figure 8B). However, actual and predicted group peak decodability numerically increased with greater numbers of electrodes (see panel of Figure 8B), as did the number of significantly decodable timepoints. We quantified these impressions via paired Wilcoxon contrasts of unbiased subject-level information timecourse peaks across electrode systems. Whereas none of the pairwise contrasts of the actual Motor network peaks were significant (p < .600 for all), contrasts of the predicted data revealed significantly greater information peaks for the 256 vs 64 systems (4.0% peak change, z=3.78, p<.001) and for the 128 vs 64 systems (3.5% peak change, z=4.33, p<.001), but not for the 256 vs 128 systems (0.5% peak change, z=0.58, p=.570). Contrasts of the subject-level prediction accuracy (i.e. predicted-to-actual temporal overlap r values) were all non-significant (p > .140 for all). Hence, while the amount of response information detectable by dynamic activity flow modeling increased with >64 electrodes, the ability of the model to capture the accompanying temporal dynamics was equivalently accurate.

The results coalesce with the source vs sensor contrasts in Figure 8A to suggest that the key findings are reliably recovered even with fewer electrodes, highlighting the robustness of our source decoding and activity flow modeling approaches across variable EEG electrode systems. However, both analyses also suggested that the strength of recovered effects increases with greater numbers of electrodes, with a prominent ‘jump’ when using greater than 64 electrodes, and more modest differences between 128 and 256 systems. This pattern has been observed in previous source modeling reports (Hassan et al., 2014; Song et al., 2015).

### Dynamic activity flow modeling is equivalently accurate across variation in model order

We also explored the effect of varying the model order parameter that determined the number of lagged predictors entered into the MVAR restFC estimation and subsequently the number of lagged activation terms used in dynamic activity flow modeling (see Figure 3). The modeling was re-run over both shorter (5 lags i.e. 25ms) and longer (15 lags i.e. 75ms) model orders. Note that the computationally intensive nature of MVAR restFC estimation meant that we had to reduce the available parameter space in a principled manner wherever possible to ensure tractability. Hence, we referenced prior relevant literature and our own work involving autoregressive modeling (see Methods for details) to select the 10 lags (50ms) model order for the main analyses. Nevertheless, these control analyses revealed that the dynamic activity flow modeling predictions were equivalently accurate across variation in model order. Figure 8C plots the model-predicted response information timecourses across the 3 model orders, which were visibly similar and yielded equivalent group peaks (see Figure legend). Paired Wilcoxon contrasts across the 3 model orders revealed no significant differences in the amount of model-predicted information captured by each model (p > .350 for contrasts of subject-level predicted timecourse peaks) nor the ability of the model to capture the morphology of the response information dynamics (p > .600 for contrasts of subject-level predicted-to-actual overlap r). Hence, the accuracy of dynamic activity flow modeling held across variation in model order.

## Discussion

The present report serves as a first demonstration of a dynamic activity flow network modeling approach capable of elucidating, in spatiotemporally resolved fashion, the network computations that give rise to multivariate information representations underlying behavior in the human brain. In developing this modeling technique, we benefited from recent advances in anatomically individualized EEG source modeling (Klamer et al., 2015; Vorwerk et al., 2018), which we combined with dynamic MVPA to decompose spatially “where” and temporally “when” task information (specifically, response and sensory information) was present across the entire brain. This extended prior applications of dynamic MVPA in sensor-level EEG, which is inherently incapable of elucidating the spatial loci of neural coding schemes and is contaminated to a greater degree (relative to source EEG) by field spread artifacts (Haufe et al., 2013; Van de Steen et al., 2019). Beyond fulfilling our need for valid spatial reconstruction, we observed generally improved information detection when sources were treated as MVPA features rather than sensors, and this held across variation in available EEG electrodes (see Figure 8). This finding should motivate increased application of source modeling in dynamic EEG/MEG decoding studies moving forwards.

Our combination of source EEG and dynamic MVPA enabled us to probe the large-scale human analogues of population coding schemes reported in animal electrophysiology research. Whilst the spatial resolution of these invasive methods remains considerably higher than our non-invasive approach, we nevertheless were able to decode with equivalent temporal resolution and superior whole-brain coverage in humans. Maximizing whole-brain coverage is non-trivial, given that regions not implanted with recording electrodes are effectively missing from descriptive maps of information content, and more critically can act as unobserved confounders (adopting the language of causal inference, Pearl and Mackenzie, 2018) in modeling how task information emerges. Our network decoding approach revealed broad representation of stimulus and response information across all major functional networks, paralleling the spatially distributed information profiles reported in animal studies (Bernardi et al., 2020; Kauvar et al., 2020; Raposo et al., 2014; Rigotti et al., 2013; Siegel et al., 2015). Despite this broad decodability, analysis of the information peak and temporal onset revealed a degree of network specialization: response information emerged earliest and maximally in the Motor network, and sensory information emerged earliest and maximally in sensory networks. Our findings align with recent electrophysiological research challenging a strictly compartmentalized or modular view of functional organization (which would not anticipate spatially distributed information), in favor of a more graded organization (Kauvar et al., 2020; Siegel et al., 2015). Whilst we are confident that our rigorous acquisition and analysis steps mitigated the influence of field spread and associated spatiotemporal leakage of neural information, future applications of our model in non-source EEG modalities (e.g. human electrocorticography, ECOG, data) would further strengthen these inferences. The simplified nature of the present sensory-motor categorization task is also worth emphasizing, and future applications of our methods to more complex tasks (and hence decoding of more complex forms of multivariate information, such as task rule/context) will be critical in interrogating the generalizability of our functional inferences. This includes our interpretation of the two-peak response decoding morphology (shown to be representationally distinct in our temporal generalization analysis, see SI Fig S3C) as reflecting early motor preparation and later motor execution/feedback processes respectively. Whilst this inference is supported by findings from animal electrophysiology and scalp EEG (Ames et al., 2019; Elsayed et al., 2016; Hanks et al., 2015; Raposo et al., 2014; Wascher and Wauschkuhn, 1996), it remains to be seen whether similar dynamic network profiles are elicited when responses are generated in different, more complex tasks.

Our dynamic activity flow network modeling analyses marked a significant advance over merely decoding task information, through formalizing a model of how that information is computed and transferred across the brain – a lacuna in the information decoding literature highlighted previously (Ito et al., 2019; Kriegeskorte and Douglas, 2018; de-Wit et al., 2016). The model accomplishes this by using connectivity estimated empirically during task-free rest to parameterize a neural network architecture, wherein these connectivity estimates weight the likelihood of task information propagating to Motor network regions from the rest of the brain. This activity flow model implemented key output innovations compared to previous versions (Cole et al., 2016b; Ito et al., 2017; Mill et al., 2020). Firstly, the model produced fully dynamic outputs of behavioral response information (with millisecond-scale resolution), unlike previous versions that predicted trial-averaged fMRI beta activations. Extending to fully dynamic predictions was facilitated by application in high temporal resolution source EEG data, and indeed the success of the model in this imaging modality (the first application in a non-fMRI modality) testifies to its general validity. A second output refinement was that predictions of future information dynamics were accomplished through a multivariate autoregression framework (past lagged task activations weighted by lagged restFC). Predicting the future information state of the brain represents a significant advance over more standard approaches of predicting held-out data at the same trial timepoint, given that this integrates a fundamental principle of causality and the direction of time (i.e. causes temporally precede effects; Granger, 1969; Lynn et al., 2021; Pearl and Mackenzie, 2018). Applying this future- predicting property to simulate the generation of response information ensured functional significance (albeit limited to this simple task context), given that this information drove subjects’ overt behavior (Williams et al., 2007; de-Wit et al., 2016).

The above output features (fully dynamic and prospective predictions) bring the activity flow framework into the realm of artificial neural network models (ANNs) that have emerged as the dominant method of simulating cognitive information in artificial intelligence (Ito et al., 2020; Yamins and DiCarlo, 2016; Yang and Wang, 2020). A critical benefit of our approach over these artificial models is its foundation in empirical data, which enabled 1) empirical estimation of network connectivity weights and 2) direct comparison of model-derived representational geometry with actual empirical data.

Regarding the first point, our ability to simulate future information dynamics in a more causally interpretable way (via past activations propagating over lagged restFC) was critically dependent on more causally interpretable estimation of restFC (Reid et al., 2019). We used an autoregressive form of multiple linear regression to estimate parameters capturing dynamic, lagged, direct and directional connectivity between brain regions. This yielded more interpretable and neuroscientifically valid connectivity weights than those estimated in ANNs via random initialization and intensive optimization. The fact that these connectivity weights generalized from predicting future activations in the rest data (implicit in how FC estimates were regularized, see Figure 3A and Methods) to making similar predictions during task performance attests both to the intrinsic (state-general) functional network architecture present at rest (Cole et al., 2014; Krienen et al., 2014), as well as the utility of autoregressive methods in capturing it (Gates et al., 2010; Liégeois et al., 2017; Mill et al., 2016). Functionally, the success of dynamic activity flow modeling provides strong evidence of the cognitive relevance of restFC (Mill et al., 2017), which yielded accurate predictions even in this difficult scenario of simulating future information dynamics in high temporal resolution data. The results therefore support the hypothesized role for restFC as a large-scale analogue of synaptic weight processes that coordinate information processing across the brain (Eichenbaum, 2018; Hebb, 1949; Wig et al., 2011).

A second benefit of our empirical modeling approach over ANNs is that it enabled direct assessment of the overlap of model-derived representations with those instantiated in the human brain. Given that the degree of computational flexibility is high even in dramatically simplified neural networks (Cybenko, 1989), this raises the possibility that accurate simulation of task information might arise from non-veridical representational codes (Kriegeskorte and Douglas, 2018; Sinz et al., 2019; de-Wit et al., 2016; Yamins and DiCarlo, 2016). To address this, we demonstrated reliable decoding of the actual data from the model-predicted representations. Predicting outputs in the same neural units as the empirical data (i.e. task activation representations in the Motor network) simplified the process of comparing the predicted and actual representational geometry, in comparison to the abstract units in ANNs. With continued advancement in computing hardware and access to large training datasets, it seems likely that ANNs will yield accurate simulations of increasingly complex neurocognitive phenomena. It is therefore vital that future ANNs aspiring towards neuroscientific insight are interrogated for their neural veridicality, rather than judging them solely on their ability to perform cognitive tasks.

Our final extension of dynamic activity flow modeling to simulate lesioning of different networks represents a particularly powerful approach (in our opinion), given that it allows for formal testing of multiple candidate neural network models in empirical data. This model comparison approach suggested a prominent role for the CCNs in representing behavioral response information: activity flow models built selectively from these three networks yielded the highest predicted response information with the earliest onsets. The high spatial and temporal precision obtained allowed us to uncover a dynamic network profile that separated the three CCNs, with early selective DAN engagement (possibly reflecting motor preparation) followed by collaborative cross-CCN engagement (possibly reflecting motor execution/feedback) in the generation of behavior.

Whilst the implied roles for the CCNs are partially consistent with previous fMRI findings linking the DAN preferentially to sensory control (Corbetta et al., 2008; Ito et al., 2017), the CON to motor control (Newbold et al., 2020b) and the FPN to flexible task control more generally (Cocuzza et al., 2020; Cole et al., 2013), separating these networks based on the precise timing and geometry of response representations was uniquely accomplished by our method’s superior temporal resolution. This separation of the CCNs by their dynamic representational profiles suggests a degree of specialization amongst these networks within their broader cognitive control functionality, which will require future applications of dynamic activity flow modeling in more complex tasks to elucidate. Future extensions of our modeling approach might also target the role of the CCNs in the *emergence* of information in the more formal sense, by disambiguating when/how new information emerges in the brain, versus charting the spread of information after its initial emergence. This will likely require refinement of dynamic activity flow modeling to integrate non-linear operators (Ito et al., 2020; Rigotti et al., 2013). Recent source modeling validations using intracranial recordings as ground truth neural generators have demonstrated reasonably accurate (∼14-24mm) EEG reconstruction of subcortical regions (Megevand et al., 2014; Seeber et al., 2019), opening up future extensions of our modeling approach at this deeper and more fine-grained spatial scale, beyond the cautious network-level focus adopted for this first demonstration. A greater emphasis on subcortical regions would also benefit from more recent functional atlases that achieve more exhaustive subcortical parcellations (Ji et al., 2019; King et al., 2019) and task paradigms that are established in engaging subcortex (e.g. medial temporal lobe activity during long-term memory tasks).

To reiterate, our modeling framework imposed certain constraints on its computations (via more causally interpretable estimation of restFC) and outputs (fully dynamic and prospective predictions of behaviorally relevant response information). This was furthered in the lesioning analysis through simulating perturbations to the system, as an analogue to classical lesioning approaches in neuropsychology (Sadaghiani et al., 2018) and disruptions to neural processing induced by brain stimulation techniques (e.g. transcranial magnetic stimulation, TMS, and transcranial alternating current stimulation, TACS). Note that the latter class of methods will undoubtedly be necessary to verify the accuracy of our simulated lesioning results, which at present merely serve to propose *candidate* network computations underlying response information flow. For example, stimulation methods can enact causal perturbations of the brain to clarify whether inhibiting (via TMS, Huang et al., 2005) or desynchronizing (via TACS, Polanía et al., 2012) CCN regions during task performance disproportionately disrupts the emergence of response information and behavior, relative to disrupting other networks.

This potential for dynamic activity flow modeling to generate predictions for stimulation interventions raises more general utility in empirically deriving individualized network models of a particular cognitive function. The efficacy in individualizing the models is demonstrated by our consistently high subject-level prediction accuracy, and previous reports highlighting the highly individualized “trait” information contained in restFC (Gordon et al., 2017; Gratton et al., 2018). The simulated lesioning approach gives a particularly clear example of how individualized network models can be manipulated/perturbed to generate insights into individuals’ neural coding profiles. Beyond providing theoretical insight into computational principles of task information, such modeling could be useful in modeling dysfunction in clinical contexts, and the successful implementation of the network models in empirical human imaging data makes this clinical translation easier than for abstract ANNs (Mill et al., 2020; Woo et al., 2017). In achieving this long-term clinical aim, we reiterate that future work will undoubtedly be necessary to test the efficacy of dynamic activity flow modeling in simulating information in more complex and naturalistic tasks, and in clinical disorders. Nevertheless, our preliminary findings attest to the potential of harnessing the causal inference tenets of this framework to build accurate, individualized models of neurocognitive function in health and disease.

## Methods

### Data and code availability

Raw and preprocessed EEG data used in the present report are accessible via the Open Science Foundation (https://osf.io/mw4k3/). The individual quantitative observations (i.e. information decoding timecourses for each subject over time) for the key Results figures (Figures 4, 5, 6 and 7) are also accessible from the same repository (subdirectory: ‘Results_figures_data’). The critical analysis code used to 1) calculate restFC via the MVAR approach, and 2) generate task activation predictions via dynamic activity flow modeling are available here https://github.com/ColeLab/DynamicSensoryMotorEGI_release.

### Participants and study design

The sample consisted of 32 healthy young adults (age mean=21.50 years, range=18-30; 14 female) out of a total of 33 recruited (one subject was excluded due to a computer malfunction aborting their session). All subjects provided informed written consent prior to participating, as per the ethical guidelines from the Rutgers University Institutional Review Board (IRB).

Each subject participated in two sessions: an MRI session (in which T1 structural MRI images were collected for EEG source modeling, in addition to T2 structural and 10 minute resting-state fMRI sequences) and an EEG session (consisting of 10 minute pre-task resting-state, ∼35 minute task and 10 minute post-task resting-state sequences). The fMRI resting-state and EEG post-task resting-state sequences were not analyzed for the present report.

### MRI and EEG data acquisition

MRI data was acquired using a 3T Siemens Trio scanner housed at the Rutgers University Brain Imaging Center (RUBIC), with a 32-channel head coil. From the full MRI session (see Participants and study design section for details), only the T1 structural images (MPRAGE, 0.8mm isotropic voxels) were used in the present report to construct realistic head models for EEG source modeling.

EEG data were acquired using the HydroCel Geodesic Sensory Net (HC-GSN) dense-array system manufactured by EGI (1000Hz sampling rate, 256 sensors, Cz reference electrode). Saline solution was used as electrolyte, and impedances across all channels were lowered to <50 kΩ at the start of the session. All resting-state and task components of the experiment were presented to subjects on a laptop running Eprime 2. To minimize fatigue and related artifacts (e.g. electromyographic, EMG, artifacts), subjects were seated in a comfortable chair and were allowed to rest freely between task blocks.

### Task design

For the EEG pre-task resting-state session (used to estimate FC weights for the dynamic activity flow model, see later Methods sections), subjects were instructed to rest whilst fixating centrally and to not fall asleep. The ensuing EEG task session consisted of a cued sensory categorization paradigm with a 2-alternative forced choice response format (2AFC; see Figure 1 for design schematic). Subjects were cued at the start of each block as to which sensory modality would be presented on the ensuing block of 10 trials, and the associated stimulus-response mappings: visual (horizontal vs vertical lines) or auditory (low vs high pitch tones). Subjects responded with the left or right index fingers across both sensory modalities, with stimulus-response mappings held constant (e.g. horizontal lines were always categorized with the left index finger). The interval between the cue and first block was randomly jittered (5.5, 6 or 6.5s in duration). On each trial, subjects had 1s to respond after stimulus onset, followed by a randomly jittered inter-trial-interval (1, 1.5 or 2s). The order of visual and auditory sensory conditions was randomized across blocks.

### Causal temporal filtering prevents circularity in dynamic activity flow modeling

Importantly, all temporal filters applied in the EEG preprocessing pipeline used a “causal” filter design (de Cheveigné and Nelken, 2019; Rousselet, 2012; Widmann and Schröger, 2012; Widmann et al., 2015). This prevented circularity from entering into our dynamic activity flow modeling analysis (see later Methods section) due to leakage of task information from the present timepoint (t0) to past lagged timepoints (t0-n lags) after filtering. Such leakage could arise from more commonly applied “non-causal” filters, wherein through both forward and backward application of the filter its output at t0 is influenced by both past (t0-n lags) and future (t0+n lags) timepoints. The backward step involving future timepoints could introduce circularity into our autoregressive dynamic activity flow modeling approach, from to-be-predicted target region activity at timepoint t0 leaking into t0-n lagged timepoints. This could lead to the target t0 timepoints being predicted to some degree by themselves. To eliminate this possibility, we used a causal filter design (one-pass, hamming-windowed, minimum phase finite impulse response (FIR); Widmann and Schröger, 2012), which only used past timepoints to calculate the output timepoints at t0. This allowed us to benefit from the critical improvements in artifact removal and signal-to-noise ratio resulting from temporal filtering, whilst not introducing circularity into the modeling analyses.

A limitation of causal filtering is the introduction of a delay in the filtered output signal relative to the input (Luck, 2005; Widmann and Schröger, 2012). Hence, we advocate caution in inferring too much into the *precise absolute* timing of the reported activation and information timecourses; rather, we focus on comparisons of *relative timing* (e.g. comparing information decoding onset across spatially distinct functional networks, which have all undergone identical causal filtering and hence are subject to equivalent output delays). Nevertheless for completeness, in the Supplementary Information (Figure S6) we provide a full set of results after application of more conventional non-causal filters during preprocessing (design: two-pass, hamming-windowed, zero-phase Butterworth (infinite impulse response, IIR)). Given that non-causal filters are designed to correct for output latency delays by backwards application of the filter, and as these filters are more commonly applied in EEG/MEG research, these supplemental results should provide some means of comparing the absolute timing of our decoding results with previous reports. Albeit it is again worth highlighting the supplementary nature of these results, given their potential for introducing leakage/circularity in the dynamic activity flow modeling results. We also encourage caution in inferring too much into absolute timing even for the non-causal filter results, as previous studies have established that these filters in practice overcorrect for the latency delay by shifting the onset of signals prior to when they actually occur (de Cheveigné and Nelken, 2019).

### EEG preprocessing

All EEG preprocessing was conducted in Matlab using the Fieldtrip toolbox (Oostenveld et al., 2011), with an identical pipeline run on the pre-task rest and task data (see Figure 2 for an overview). Notch filtering was firstly applied to the continuous data to remove line noise (cutoffs at 60Hz, and higher harmonics 120Hz and 180Hz), followed by high-pass filtering (1Hz cutoff). For all filters, a causal filter design was used (as detailed in the preceding section) and filter order was optimized in a lower-level Fieldtrip function to ensure the highest slope for the cutoff frequencies. Noisy sensors were then identified via an automated procedure, wherein all sensors with a zscored mean absolute amplitude (across all timepoints) greater than 3 were removed. The continuous data was segmented into trials: -0.5 to 1.5s for the task data around trial stimulus onset, and into 20s non-overlapping “pseudotrials” for the rest data. Noisy trials/pseudotrials were identified via a similar automated procedure: all trials with a zscored mean absolute amplitude (averaged across all trial timepoints and sensors) greater than 3 were removed.

Temporal independent components analysis (ICA; Jung et al., 2001) was used to identify eye blink and eye movement artifacts. Extended ICA was run on each subject’s task and rest data using the “binica” method, accessible in Fieldtrip via an EEGLAB plugin (Delorme and Makeig, 2004). After estimating the component scores, a semi-automated procedure was used to identify components capturing ocular artifacts. Two vertical electro-oculogram channels (EOG) located above and below the right eye were subtracted from each other to provide a timecourse capturing blink artifacts with a high signal-to-noise ratio (Chaumon et al., 2015). Similarly, two horizontal EOG channels located adjacent to the right and left eyes were subtracted from each other for the eye movement/saccade timecourse. These vertical and horizontal EOG signals were used as “artifact templates” that were separately correlated with each ICA component score’s timecourse (following a similar approach in Pontifex et al., 2017), with likely artifact-related components identified as those with a zscored absolute Pearson r value > 3. These candidate artifact components were finally accepted/rejected after visual inspection. The ICA-cleaned data were then low-pass filtered (50Hz cutoff) to reduce high frequency artifacts (e.g. EMG artifacts which peak in the 50-100 Hz range; Goncharova et al., 2003), baseline corrected (treating -0.5 to 0 as the baseline for the task trial data, and the mean over each entire pseudotrial as the rest baseline) and re-referenced to the common sensor average.

### EEG source modeling

The preprocessed task and rest EEG data then underwent source modeling to recover the underlying neural source activations from the sensor signals (Figure 2). Source modeling was critical for analyses requiring valid recovery of spatial signatures of task information representation, versus relying on sensor-level analyses that are inherently spatially ambiguous (Haufe et al., 2013; Pernet et al., 2020; Reid et al., 2019; Van de Steen et al., 2019). This endeavor was helped by our use of a dense-array EEG system (256 sensors), given that such a high number of sensors has been previously shown to improve EEG source localization accuracy to match the millimeter precision observed with MEG (Klamer et al., 2015; Lascano et al., 2014; Michel and Brunet, 2019; Mill et al., 2016; Seeber et al., 2019).

Realistic, individualized head model construction began by segmenting the anatomical T1 MRI images for each subject into five skull tissue types (gray matter, white matter, cerebrospinal fluid, skull and scalp). This enabled finite element modeling (FEM; Vorwerk et al., 2018) of the distortion of electric dipole fields generated by neural activity as they propagate through skull tissues towards the scalp sensors. The FEM head model was then aligned to the dense-array EEG sensors based on common fiducial landmarks. Source region locations were specified in MNI space based on the whole-brain functional atlas described by Power et al (Power et al., 2011), which provides independent functional network affiliations for 264 regions. These network affiliations were identified via clustering analyses applied to task and resting-state fMRI data in the original paper, and have subsequently been verified via alternative (Gordon et al., 2016) and multi-modal (Glasser et al., 2016; Ji et al., 2019) human parcellation approaches. Convergent network profiles have also been observed in animals (Stafford et al., 2014; Wang et al., 2013), highlighting the general validity of this functional atlas. The source regions were aligned linearly with the head model and sensors. The full forward model was then estimated via normalized leadfields describing electric field propagation from the neural sources through skull tissues to the recorded EEG electrodes.

Beamforming in the time domain via the linearly constrained minimum variance approach (LCMV; Van Veen et al., 1997) was used to invert the forward model and reconstruct the activation timeseries for each source location. We opted for beamformer-based source modeling as this has been shown to mitigate artifactual field spread influences in source connectivity analyses, compared to alternative minimum norm estimate (MNE) approaches (Schoffelen and Gross, 2009). The orientation of the reconstructed source timeseries was constrained to the direction yielding maximum power (based on singular value decomposition of the 3-dimensional xyz dipole moments). This entire source modeling procedure was applied separately to the task and rest EEG data, reconstructing the relevant activation timeseries for each session from the same 264 spatial locations.

### Dynamic multivariate pattern analysis (MVPA)

The source-modeled task EEG data were then submitted to dynamic multivariate pattern analysis (MVPA) to decode task information with millisecond-scale temporal resolution. The analyses focused primarily on decoding behavioral response information (left-vs right-handed responding, Figure 1) from response-locked data, but decoding of sensory information (visual vs auditory stimulation) from stimulus-locked data was also conducted as a supplementary analysis (see SI, Figure S1). The features used for dynamic MVPA were separately “SensorAll” (timeseries from all 256 EEG channels), “SourceAll” (timeseries from all 264 source regions localized from the Power atlas) or “network sources”. For the latter, dynamic MVPA was performed for each of the 11 Power atlas networks^2^ separately (treating within-network source regions as features), yielding 11 information decoding timecourses.

The steps adopted were the same across these feature sets. For response information decoding, the trial data were first rearranged from stimulus-to response-locked (segmented -.45 to .45 seconds around response commission), using the trial reaction times. The data were then downsampled from 1000Hz to 200Hz (by taking every 5th sample sequentially) to boost signal-to-noise for the classifications (as recommended in Grootswagers et al., 2017). Incorrect trials were removed and the remaining trials arranged into the two conditions of interest: correct left and correct right response trials, collapsed across visual and auditory stimulation conditions. For each response condition, trials were averaged into smaller “subtrial” sets to boost signal-to-noise for the classifications (Grootswagers et al., 2017). Specifically, the ∼120 trials in each condition were averaged over non-overlapping sets of 14 trials to yield ∼8 independent subtrials per condition, per subject (‘∼’ reflects variable removal of noisy and incorrect trials across subjects during preprocessing). The trial indices going into these subtrial averages were randomized over 10 iterations (whilst preserving non-overlap within each iteration), meaning that the entire decoding analysis was run 10 times per subject to ensure robustness of the resulting information timecourse.

Within each of the 10 subtrial averaging iterations, linear support vector machine (SVM) classifiers were trained separately at each trial timepoint to distinguish between left-and right-handed responses, as implemented by libSVM (Chang and Lin, 2011) and the cosmoMVPA toolbox (Oosterhof et al., 2016). Ten-fold crossvalidation was used, in which trials were pseudorandomly split into 80% training and 20% testing sets in each of 10 folds, with the constraint that no unique combination of train and test trial indices were repeated across folds. Note that the number of training trials were rounded up (e.g. 20% of 8 available trials=1.6=2 training trials per condition) and the number of testing trials were rounded down (e.g. 80% of 8 trials=6.4=6 testing trials per condition), following the default behavior of cosmoMVPA. Note also that the number of train/test trials selected from each response condition was equated on each fold to prevent imbalanced trial counts from influencing the classifiers (Poldrack et al., 2019). The accuracy (%) in classifying between left- and right-handed responses was averaged over folds, with this classification process repeated for each individual timepoint to yield a decoding timecourse capturing response information dynamics for that subject. The final decoding timecourse for each subject was taken as the average across the 10 subtrial averaging iterations. To summarize, the subtrial averaging procedure was looped 10 times (with randomized initial trial indices), with 10-fold cross-validation then initiated separately within each subtrial loop, and the decoding results averaged across the cross-validation folds and subtrial loops to derive the information timecourse for that subject.

Statistical significance was assessed via a non-parametric approach treating subjects as a random effect, wherein the classification accuracies at each timepoint were contrasted against chance classification (50%) via the Wilcoxon signrank test (Grootswagers et al., 2017). The resulting p values were Bonferroni-corrected for multiple comparisons across timepoints. For analyses contrasting decoding peaks via paired Wilcoxon signrank tests, peak classification accuracies were extracted from each subject’s decoding timecourse from the relevant group time-to-peak (see Results for details).

The Supplementary Information provides a number of other network decoding analyses, primarily conducted to validate the spatial accuracy of source modeling: decoding of sensory information (visual vs auditory stimulation) from stimulus-locked data (Figure S1 panels B-D); decoding of stimulus information (visual: horizontal vs vertical lines; auditory: low vs high pitch tones) from stimulus-locked data (Figure S2); decoding of response information from stimulus-locked data (Figure S3 panels A-B). We performed a supplementary temporal generalization analysis (King and Dehaene, 2014) to probe whether the multivariate codes underlying the two peak deflections in the response decoding timecourse were representationally distinct (SI, Figure S3C). We also conducted network decoding of response information using an alternative method that confined MVPA to more “unique” signals for each network (SI, Figure S4), which revealed a highly similar pattern of results.

### Multivariate autoregressive (MVAR) estimation of resting-state functional connectivity: rationale

The source-modeled rest EEG data were used to estimate the lagged functional connectivity weights (Figure 3A) that parameterized the dynamic activity flow model (Figure 3B). Note that we only used the pre-task rest phase here (see Participants and study design section), given the potential for influences of the task on FC in the post-task rest phase. These could encompass second-order connectivity changes induced by the task (Muraskin et al., 2016) or first-order coactivation effects from cognitive replay of the task (Cole et al., 2019), which could have introduced circularity in predicting activation/information in the earlier task phase.

Our motivations for using a multivariate autoregressive (MVAR) FC approach stemmed from the overarching aim of imposing certain constraints on the subsequent dynamic activity flow modeling, taking inspiration from concepts in the causal inference (Pearl and Mackenzie, 2018) and causal connectivity (Mumford and Ramsey, 2014; Ramsey et al., 2010) literatures. Note that by imposing these modeling constraints we do not claim to have effectively solved the problem of identifying causal relationships between brain regions. Rather, we took these steps as a principled means to reduce the available space of inquiry and improve (in a relative sense) the likelihood of identifying true causal relationships in the brain via our adopted MVAR FC approach compared to other less constrained methods (e.g. Pearson correlation, univariate/bivariate autoregressive modeling). We have delved into these concepts in detail in prior theoretical work (Mill et al., 2017; Reid et al., 2019).

For the first of these principled reduction steps, MVAR permits estimation of dynamic aspects of FC through the inclusion of predictors over temporally lagged timepoints (Liégeois et al., 2017; Mitra and Raichle, 2016), in contradistinction from static FC methods (e.g. Pearson correlation or multiple linear regression estimated over an entire rest session). Secondly, our MVAR approach allows for separation of contemporaneous (connectivity of region A at t0 with region B at t0) and lagged (connectivity of region A at t0-1 lag with region B at t0) contributions to FC (Gates et al., 2010; Kim et al., 2007). This is unlike typical sliding window dynamic FC approaches, which assess fluctuations in contemporaneous influences over discrete time windows (Hutchison et al., 2013), and have been criticized for potentially capturing sampling variability (i.e. noisy estimates of static FC) rather than true dynamic interactions (Laumann et al., 2016; Liégeois et al., 2017). Our emphasis on the “lagged” nature of our approach is predicated on modeling both contemporaneous and lagged terms in the MVAR model so as to estimate the true lagged FC effects. This differentiates our approach from alternative autoregressive methods that also target dynamic FC influences by modeling lags, but do so without accounting for contemporaneous ones (leading to errors in lag estimation, as previously reported; Gates et al., 2010). Estimation of dynamic FC with clear separation of lagged influences was essential to our goal of predicting *future* activation/information states via dynamic activity flow modeling.

Thirdly, our MVAR approach fitted all spatial (“source” predictor regions) and temporal (t0-n lags) terms simultaneously via multiple linear regression models (Figure 3A), meaning that the approach is fully multivariate in both the temporal and spatial domains. This reduces the impact of indirect spatial or temporal influences from serving as unobserved confounders on the FC weights, which would likely arise to a greater extent if alternative bivariate FC estimation methods were used (Pearl and Mackenzie, 2018; Reid et al., 2019; Sanchez-Romero and Cole, 2020), or if alternative neuroimaging modalities were used (e.g. the poor temporal resolution of fMRI allowing for temporal confounding, and the poor whole-brain coverage of animal electrophysiology allowing for spatial confounding). Whereas the risk of unobserved confounders still persists (Stevenson, 2018), as do related problems introduced by source EEG data (e.g. the coarse spatial scale of EEG, field spread artifacts), we have at least reduced this risk by adopting a multivariate rather than univariate approach to restFC estimation and network modeling. Fourthly, our exclusive focus on the lagged MVAR estimates for dynamic activity flow modeling (see later Methods section for details) imparted directionality to the approach, given that past t0-n lagged terms predicted future target t0 terms (Figure 3B). This autoregressive form of directionality is the basis of Granger Causality methods (Geweke, 1982; Granger, 1969), which have been fruitfully applied to both fMRI and source modeled MEG/EEG data (Cheung et al., 2010; Mill et al., 2016; Roebroeck et al., 2011).

To summarize, each of the four terms we use to describe our approach is intended to clarify a principled constraint versus alternative approaches: dynamic (vs static), lagged (estimating both lagged and contemporaneous effects, vs the lags alone), direct (multivariate in spatial and time domains, vs univariate) and directional (modeling future activity from the past lagged terms vs modeling them from lagged and contemporaneous ones). MVAR yielded FC estimates that hence increased the overall validity - according to extant causal inference/connectivity frameworks (Mill et al., 2017; Reid et al., 2019) - of this version of activity flow modeling beyond previous versions.

### MVAR estimation of resting-state functional connectivity: specific approach

The rest data (already segmented into 20s non-overlapping pseudotrials^3^, see EEG preprocessing section) were firstly downsampled from 1000Hz to 200Hz. This matched the downsampling applied to the task data and reduced computation time for the intensive MVAR procedure. Consistent with our focus on predicting activation/information states in the Motor network via dynamic activity flow modeling (see next section), the multiple linear regression models comprising the MVAR step treated the 35 Motor network regions as to-be-predicted (target, j in Figure 3A) regions, and the remaining network regions as predictor (source, i) regions. Timeseries for all regions were firstly zscored within each pseudotrial. A model order of 10 lags was selected (i.e. t0-10 timepoints extending 50ms into the past of each target t0 timepoint), based on evidence from invasive electrophysiology suggesting 50-100ms as a reasonable timescale over which task information fluctuates (Crowe et al., 2013; Murray et al., 2014; Siegel et al., 2015), and our previous work directly involving autoregressive modeling in EEG/MEG data that used these parameters (Cole et al., 2010; Mill et al., 2016). We opted for the lower bound of this range (50ms) given the computationally intensive nature of regularized MVAR restFC estimation (3-5 days runtime per subject), which scaled with the number of predictors entered in the PCA autoregression model and hence increased with larger model orders. As a control analysis, we also present dynamic activity flow model accuracy results across variation in model order (25ms and 75ms; see Results, Figure 8C).

When selecting MVAR predictor terms, our general approach was to estimate all possible spatial and temporal terms to eliminate indirect influences arising from unobserved confounders, with more selective use of predictors in the later dynamic activity flow modeling step. Hence, all non-target Motor network regions were included in the MVAR source set, despite these regions being excluded from dynamic activity flow (to focus the model on long-distance connections, see next section).

Similarly, both contemporaneous (t0) and lagged (t0-10 lags) terms were included for source predictor regions during the MVAR step, despite basing the dynamic activity flow modeling solely on the lagged source terms (to preserve our aim of predicting *future* activation/information states). This follows previous simulation work demonstrating that inclusion of contemporaneous terms improves the precision of lagged estimates, versus fitting the lagged terms alone (Gates et al., 2010). The MVAR models for each target region also included a set of autoregressive lagged terms (over t0-10, excluding t0 to prevent circularity) that estimated serial autocorrelation or self-coupling as per common guidelines (Gates et al., 2010; Geweke, 1982; Liégeois et al., 2017).

The MVAR FC weights for the input spatial and temporal terms were estimated via a regularized form of multiple linear regression, involving principal component analysis (PCA; Figure 3A). We have applied a non-autoregressive variant of this FC approach recently (Mill et al., 2020), and the regularization here was similarly focused on identifying the number of predictor PCs (nPCs) included in each MVAR regression that would reduce overfitting to noise. The approach employs cross-validation to identify the optimal number of principal components to include in regression models estimated for each subject, so as to regularize the output FC beta coefficients and minimize overfitting. To ensure tractability during this computationally intensive procedure, nPCs were varied from 1 to the max number of PCs (i.e. the rank of the predictors, ∼2700) in increments of 540.

Hence, for a given Motor target region on a given rest pseudotrial, PCA was run on the accompanying predictor matrix (263 contemporaneous source + 263*10 lagged source + 10 lagged self-coupling terms=2903 predictors, fit to each 20s pseudotrial of 4000 observations). The nPCs retained were varied as per the regularization (1:540:max), with the target region’s timeseries then regressed onto those selected nPC scores. The resulting beta weights were transformed back to the original predictor space, and then applied to predict the target timeseries for the next pseudotrial. The mean squared error (MSE) between the actual and predicted next-pseudotrial timeseries was stored, with this process repeated for all trials in sequence (i.e. train on pseudotrial1, test on pseudotrial2; train on pseudotrial2, test on pseudotrial3 etc). Note that this corresponds to a “sliding window” cross-validation scheme that preserves the serial ordering of the input timeseries during model training (unlike more general k-fold cross-validation schemes) to theoretically improve generalization (Cerqueira et al., 2019). The full regularization scheme hence generated an MSE output variable with dimensions nPCs x target regions x test pseudotrials, with the optimal nPCs finally selected as that yielding the minimal MSE (averaged across target regions and test pseudotrials) for that subject. This optimal nPC value was used to estimate the final MVAR restFC weights for all Motor network targets across all pseudotrials for that subject, with the cross-pseudotrial average restFC matrix used in the next dynamic activity flow modeling step.

### Dynamic activity flow modeling

We applied dynamic activity flow modeling to capture the emergence of future response information representations in the Motor network. The model is depicted graphically in Figure 3B, and summarized in the following equation:

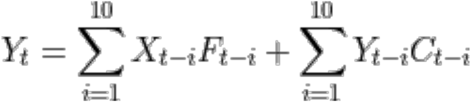

Where t represents a future to-be-predicted task trial timepoint, i the set of lagged timepoints extending 10 timepoints into the past, Y the to-be-predicted regional activation (and lagged autoregressive terms on the right of the equation), X represents the lagged source (non-Motor network) predictor regional activations, F represents the lagged source predictor restFC terms, and C represents the lagged autoregressive/self-coupling restFC terms.

The present model built on previous versions (Cole et al., 2016b; Ito et al., 2017; Mill et al., 2020) by including stronger modeling constraints. As introduced in the previous MVAR FC estimation sections, these constraints ensured that predictions of future response information states were made from activity flowing over functional network connections in a dynamic, lagged, direct and directional fashion. This more principled grounding was strengthened by focusing the modeling on response information representations, given that these underlie generation of overt behavior and hence likely mark a causal end-point of cognitive task processing (Williams et al., 2007; de-Wit et al., 2016). This justified imposing directionality from t0-n lagged non-Motor sources→t0 Motor targets in the model (Figure 3B). Such temporal directionality would be harder to ascribe in dynamic activity flow models of sensory information in the Visual network (following up on the sensory decoding results, see SI Figure S1), given greater alternation between feedforward (activity flow from lagged non-Visual sources→Visual targets) and feedback cycles (lagged Visual sources→non-Visual targets; Gwilliams and King, 2020).

To reiterate, certain predictor terms that were modeled in the MVAR restFC step were excluded from the ensuing dynamic activity flow modeling: non-target Motor regions and contemporaneous (t0) source terms. These exclusions focused the predictions on spatially long-range and temporally in-the-future activity flow processes respectively. Lagged autoregressive terms were included to account for local self-coupling processes that likely contribute to dynamic activity flow. The source-modeled task EEG data were firstly downsampled to 200Hz. Predicted activations for each target Motor network region (j in Figure 3B), for each task trial, for each timepoint (t0), were then generated from the dot product of lagged task activations (for sources and self-coupling terms, over the same model order of t0-10 samples) and lagged rest MVAR restFC weights (capturing FC *from* the sources and self-coupling terms *to* the targets). This generated a matrix of predicted Motor region task activations with the same dimensions (Motor regions x task trials x timepoints) as the actual task activations used for the previous decoding analyses. This predicted activation matrix was the basis of subsequent assessments of model accuracy: response information decoded via dynamic MVPA, motor ERP analyses and representational overlap (see ensuing sections).

Some further features of the model that aimed to rigorously rule out circularity are worth highlighting. To reiterate, causal filtering was used during preprocessing to prevent introduction of circularity due to temporal leakage (i.e. task activation from targets at t0 leaking to lagged predictors after filtering, see earlier Methods section). Note also that the MVAR FC weights were derived from a resting-state session that was entirely separate from the task activation data, which prevented circularity in estimating FC from the same task session as the to-be-predicted data (due to first-order task coactivation effects contaminating the FC weights; Cole et al., 2019). We also implemented multiple strategies to mitigate circularity introduced by EEG field spread artifacts, which could lead to target t0 activations spreading to spatially proximal predictor regions. Firstly, we employed beamformer source modeling which has been shown to minimize such artifacts compared to alternative methods (see EEG source modeling section; Schoffelen and Gross, 2009). Our use of the Power functional atlas (Power et al., 2011) to identify source regions also helped with this goal, given that each region is spaced at least 10mm apart from each other and hence avoids the most severe field spread arising between highly proximal regions (Schoffelen and Gross, 2009). Our exclusion of the contemporaneous (t0) source terms from the activity flow step also mitigates field spread, as these artifacts are instantaneous and hence unlikely to influence lagged terms (Nolte et al., 2004; Stinstra and Peters, 1998). Our careful use of causal filters as highlighted above eliminated any residual possibility of field spread-related signals leaking from a target timepoint t0 to lagged timepoints in the past. As a final rigorous control, we also regressed out the task timeseries for a given target (t0) from all predictor source regions (at the same t0), prior to rearranging them into lagged predictors for dynamic activity flow modeling. This regression step was run just prior to all dynamic activity flow modeling analyses presented in the main manuscript, and effectively removed all contemporaneous influences (including those resulting from field spread and temporal leakage of information) on the model outputs. The supplement describes how this step actually numerically improved the model prediction accuracy (versus omission of this step; see SI, Figure S7).

### Assessment of dynamic activity flow model accuracy: dynamic MVPA

The accuracy of dynamic activity flow modeling was assessed in a number of ways (Figure 3B). Our primary approach was to apply the identical dynamic MVPA procedure to the model-predicted Motor region activation timeseries as was applied to the actual data (see Dynamic MVPA section). This yielded a predicted response information timecourse, for which significant decodability at each timepoint was assessed via Wilcoxon signrank against 50% chance (Bonferroni-corrected across multiple timepoint comparisons, as before). The recovery of significantly decodable timepoints in the predicted timecourse would provide evidence that our dynamic activity flow modeling approach accurately captured the emergence of future response information. The success of the model in capturing the temporal morphology of the response information timecourse was also quantified via Pearson correlation of the predicted and actual timecourses. This was computed both after averaging the timecourses across subjects (group-level overlap), as well as with a random effects approach that correlated the predicted and actual timecourses for each subject and contrasted this r value (after Fisher-z transform) against 0 via one-sample ttest (subject-level overlap).

We also report the coefficient of determination (R^2^) computed using the sum of squares formulation (Poldrack et al., 2019) for all dynamic activity flow models. This metric permits assessment of scaled prediction accuracy (unlike Pearson r), as well as providing insight into whether the model predictions outperform those based on the average actual data. To quantify whether R^2^ was reliably greater than 0 (indicating that dynamic activity flow modeling outperformed the average model), we adapted the random effects approach used for the Pearson r metrics: computing R^2^ for each subjects’ predicted/actual data, and contrasting at the group level against 0 via one-sample ttest. For the dynamic MVPA analysis, we also conducted a permutation testing analysis to further highlight the prediction accuracy of our model. This is described in the Results.

### Assessment of dynamic activity flow model accuracy: motor event-related potential (ERP)

We also examined the model’s accuracy in capturing the activations in Motor network regions that underpinned the dynamic MVPA analyses. This targeted recovery of the well-established motor ERP termed the lateralized readiness potential (Cheyne et al., 2006; Deecke et al., 1976; Kutas and Donchin, 1980). This reflects greater activation around response commission in the Motor network hemisphere contralateral to the response hand. For each subject, the model-predicted trial activations were arranged into “contralateral” (left-response for right hemisphere regions and right-response for left hemisphere regions) and “ipsilateral” (left-response for left hemisphere and right-response for right hemisphere) conditions, and averaged across regions and trials. The difference wave for the resulting contralateral minus ipsilateral waveforms hence indexed recovery of the motor ERP for each subject, with this entire process applied separately to the model-predicted and actual data. In both predicted/actual cases, significance was assessed by contrasting the subject motor ERP fluctuations at each timepoint against 0 via Wilcoxon signrank test, with Bonferroni correction for multiple timepoint comparisons.

The accuracy of the model in predicting the raw task activation effects was assessed firstly as the ability to recover significant activations in the predicted motor ERP waveform. We also quantified the model’s ability to recover the morphology of the motor ERP by correlating the predicted and actual difference waves, at both the group-and subject-levels. Whilst this constituted an assessment of “temporal overlap”, we were also able to probe predicted-to-actual overlap in the spatial domain as the ERP analyses did not aggregate information across spatially distinct regions (i.e were spatially univariate unlike the dynamic MVPA analyses). The subject motor ERPs were averaged over significant epochs identified in the group actual waveform, separately for each Motor region, and separately for the predicted and actual data. The predicted and actual activation vectors were then correlated at the group- and subject-levels to interrogate how well the dynamic activity flow model captured the spatial pattern of Motor network activations. To ensure that spatial overlap was not critically dependent on the selected significant epoch, we also assessed group- and subject-level overlap between predicted and actual activation vectors that concatenated all Motor regions and all trial timepoints (i.e. capturing the degree of “spatiotemporal overlap” across the entire trial timecourse).

### Assessment of dynamic activity flow model accuracy: representational overlap

We also probed the extent to which the multivariate representations underlying the accurate decoding of response information in the activity flow-predicted data overlapped with the representations in the actual data. Such direct “representational overlap” between the predicted and actual data would increase confidence that dynamic activity flow modeling is capturing how the brain veridically represents response information. To address this, we tested whether response information in the actual data could be decoded from multivariate representations trained in the predicted data. This approach of training in the predicted and testing in the actual data is in contrast to that adopted in the main decoding analysis (Figure 6A), wherein training and testing was performed in the predicted data. We employed a method inspired by representational similarity analysis (Diedrichsen and Kriegeskorte, 2017; Ito et al., 2017; Mur et al., 2009), which was capable of more fine-grained assessment of the similarity in representational geometry (by computing continuous Pearson r similarity values as the measure of classification accuracy) than the SVM classification approach used for the main dynamic MVPA analyses (which output categorical decisions to assess accuracy).

Predicted task activations for Motor regions were generated via dynamic activity flow modeling as described above, which were then averaged into subtrials separately for the two response conditions (correct left and correct right) and assigned into training and test sets via 10-fold cross-validation (as with the dynamic MVPA analyses). Representational templates were created at each timepoint for each response condition by averaging Motor region activations over relevant trial indices in the *predicted* data. These timepoint-by-timepoint predicted representations were then applied to decode the *actual* Motor activations in the held-out test trials. For each timepoint in each test trial, Pearson correlations were computed between the actual activation vector and both the “correct” predicted condition template (e.g. left response template correlated with actual left response test trial) and the “incorrect” predicted condition template (e.g. right response template correlated with actual left response test trial). Hence, the difference value for correct r minus incorrect r provided an estimate of response information decodability at each timepoint in the actual test data (with values > 0 denoting presence of information). Iterating this process over all timepoints, testing folds and subtrial iterations for each subject generated a timecourse capturing the degree of overlap between the predicted and actual response information representations.

Statistical significance of the representational timecourse was again examined via Wilcoxon signrank tests against 0, with Bonferroni correction across multiple timepoint comparisons. Recovery of significantly decodable timepoints in this “TrainPred-TestActual” timecourse provided evidence that the dynamic activity flow model accurately captured representations underlying future information decoding in the actual data. For comparison, we repeated the same representational similarity approach with trained condition templates computed in the actual data, thereby generating a “TrainActual-TestActual” timecourse. Correlating the TrainPred-TestActual and TrainActual-TestActual timecourses (at both the group- and subject-levels) hence clarified how well the activity flow model captured dynamics in representational geometry over the entire trial.

### Network lesioning extension of dynamic activity flow modeling

To extend the dynamic activity flow model towards insight into principles of functional brain organization, we applied a modified “network lesioning” variant of the model. This followed a similar approach to generating activity flow-predicted response information timecourses for the Motor network via dynamic MVPA, as described above. The critical difference here was that information timecourses were predicted via separate activity flow models that selectively “lesioned” all except one of the 11 functional networks from the Power atlas. Hence, only regions within one specific network were included in the predictor source set (Figure 3B) for each dynamic activity flow model, with lagged self-coupling/autoregressive terms also excluded. This yielded 10 timecourses capturing the unique contributions to response information decoding made by each individual network.

Comparing peak decodability across network models hence provided more computational insight into which networks were dominant drivers of response information. Such comparisons were made via visual inspection of the 10 lesioned information timecourses, as well as by contrasting decoding peaks (extracted for each subject from the group time-to-peak across networks) via paired Wilcoxon signrank tests (with FDR correction for multiple pairwise network comparisons). As an alternative to contrasting peak decoding accuracies, we also contrasted the subject-level prediction accuracy for each network model (i.e. Pearson r for each network-predicted timecourse with the actual Motor network timecourse) via paired Wilcoxon signrank tests (with FDR correction for multiple pairwise network comparisons). This captured how well each lesioned network model predicted the morphology of the actual response information timecourse (SI, Figure S8).

## Acknowledgements

The authors acknowledge support by the US National Institutes of Health under awards R01 AG055556 and R01 MH109520 to MWC. The authors acknowledge the Office of Advanced Research Computing (OARC) at Rutgers, The State University of New Jersey for providing access to the Amarel cluster and associated research computing resources that have contributed to the results reported here. The authors thank Dr Bart Krekelberg for providing access to the EEG equipment used to collect data for this report, which was purchased with funding secured by Dr Krekelberg from the US National Science Foundation (project BCS-1530930). We also thank Jasmine Siegel for assisting during installing and collecting data using this equipment. The content is solely the responsibility of the authors and does not necessarily represent the official views of any of the funding agencies.

## Author contributions

RDM and MWC designed the task paradigm and analysis approach. RDM, JLH, ECW, NL and RHC were responsible for data collection. RDM performed the analysis under the supervision of MWC, with contributions from JLH, ECW and NL. RDM and MWC wrote the manuscript, with feedback received from all other authors.

## Footnotes

1 Results of the PCA regression regularization (achieved via cross-validated optimization): cross-subject mean squared error (MSE) in predicting future resting-state activation timeseries=0.006 (std=0.003); optimal nPCs=540 (59.4% of subjects) or 1080 (40.6%).

2 For all analyses targeting inferences at the network level (i.e. the network decoding and all dynamic activity flow modeling analyses), we excluded all regions from the “Uncertain” and “Retrieval” Power atlas networks, given that these networks have not been reliably recovered in more recent parcellation schemes (e.g. Ji et al., 2019). For similar reasons, we collapsed the original “Somatomotor-hand” and “Somatomotor-face” (depicted in orange in Figure 2, panel iii) networks into a single “Motor” network definition.

3 We selected 20s as the length of each pseudotrial to capture the lower bound of the low frequency fluctuations considered to contribute to resting-state FC (i.e. 0.1-1Hz, with 20s allowing 2 cycles of 10s; Kucyi et al., 2018; Vincent et al., 2007), as well as contributions from higher frequencies.

