## Supplementary Information for "Network modeling of dynamic brain interactions predicts emergence of neural information that supports human cognitive behavior"

**1. Sensory information decoding results.**
Figure S1 plots the results of applying dynamic MVPA to decode sensory information (Visual versus Auditory task conditions, see Figure 1) from stimulus-locked data. Figure S1A plots the comparison of the SourceAll versus SensorAll sensory decoding timecourses, which accompany the same comparison for response decoding in Figure 4. Multiple significantly decodable timecourses were observed in both timecourses, which followed a sensible morphology: decodability was at baseline prior to stimulus onset and rose to peak between 100-200ms. The group sensory decoding peak was numerically higher for the SourceAll versus the SensorAll timecourse, paralleling the direction of the response decoding comparisons in Figure 4. Formal contrasts of the decoding peaks via paired Wilcoxon signrank tests did not reach significance, across variation in how peaks were defined: i) from the SensorAll/SourceAll group time-to-peak (both 0.175s; 1.2% peak change; $z=1.01$, $p=.312$), ii) from the peak of each individual subject's decoding timecourse (unbiased by the group results; 0.02% peak change; $z=0.12$, $p=.905$). Hence, the results revealed numerical albeit non-significant improvements in overall decodability of sensory information for the source-modeled dynamic MVPA approach.

Figure S1B plots network-level decoding of sensory information, with Figure S1C and S1D ranking the cross-network decoding peaks extracted from two group peak timepoints (at 0.175 and 0.265s respectively). The network decoding timecourses again revealed content-appropriate specialization: sensory information peaked highest in the Visual network, with an early onset. Similar to its involvement in the likely sensory-related peak1 of the response decoding timecourse (see Figure 5), the DAN was also prominently involved in representing sensory information here, yielding the third-highest overall peak and earliest onset. Pairwise contrasts of peak decodability (accompanying Figures S1C and S1D) confirmed this preferential engagement of the DAN, which decoded significantly higher than the CON at the sensory peak1, and significantly higher than both the CON and FPN at peak2 ($p<.05$ via paired Wilcoxon, FDR-corrected for multiple pairwise contrasts). Overall, network decoding of sensory information highlighted a more selective engagement of the DAN amongst the CCNs, in contrast to the more collaborative CCN engagement observed during motor execution/feedback (see Figure 5D and 5E).

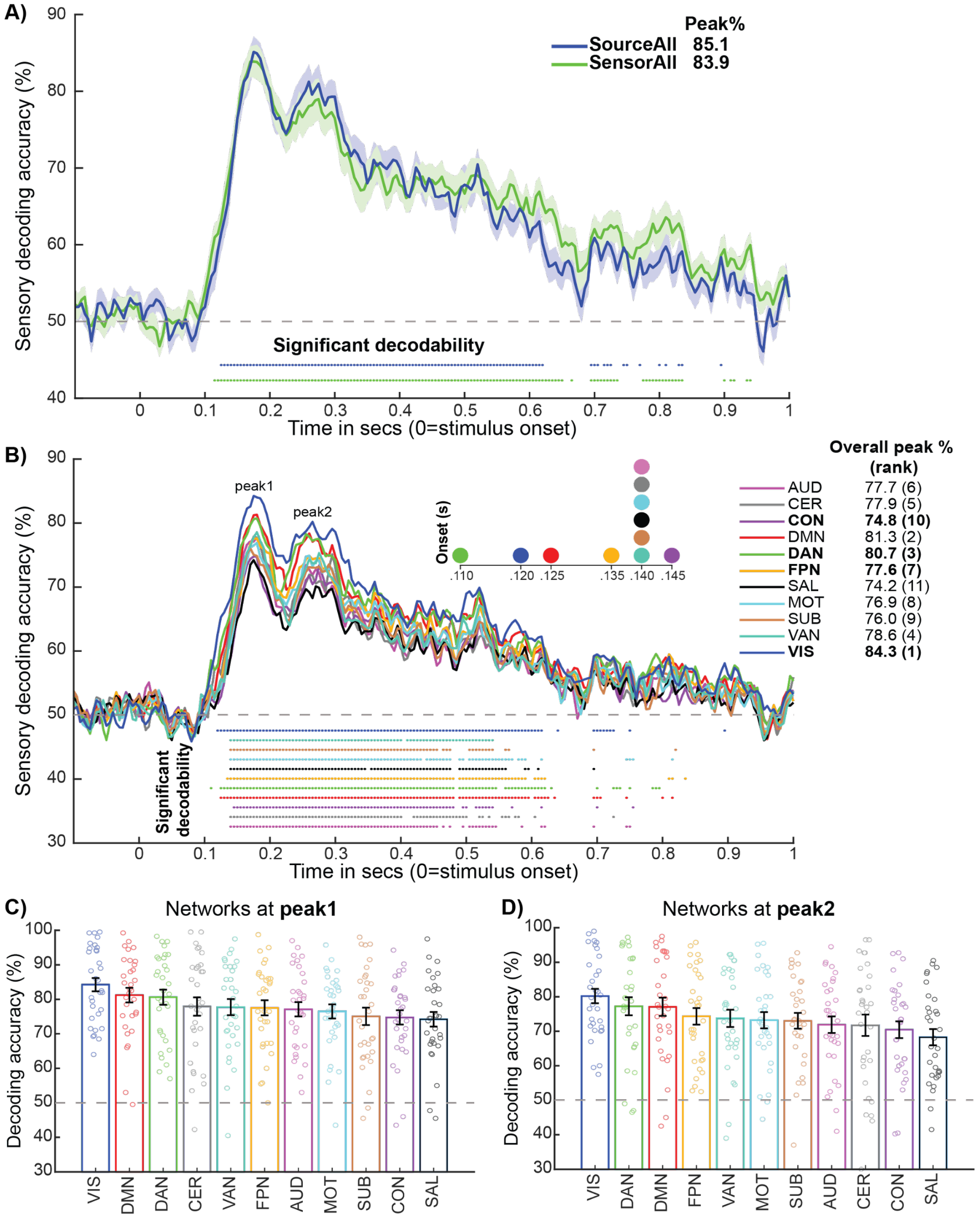


**Figure S1. Sensory information decoding results. A)** Comparison of SourceAll versus SensorAll group decoding timecourses. **B)** Network level decoding of sensory information, with ranked networks peaks extracted from **C)** peak1 (0.175s) and from **D)** peak2 (0.265s). Plotting conventions are the same as those in Figure 5 in the main manuscript.

### 2. Content-appropriate decoding of auditory and visual stimulus categories further highlights source modeling precision.

The spatial accuracy of our source modeling approach was demonstrated by response and sensory information peaking in their content-appropriate Motor (Figure 5A) and Visual (Figure S1A) networks respectively. To further highlight our source modeling accuracy, we also decoded stimulus categories within each sensory modality: horizontal versus vertical lines for the visual condition; low versus high pitch sounds for the auditory condition. Given the one-to-one assignment of each condition to a response hand (e.g. horizontal/vertical lines were always identified with left-/right-handed responses), we restricted this analysis to an early epoch after stimulus onset (-0.1 to 0.3s) to prevent contamination by later left versus right hand response information decoding. The results revealed that visual stimulus information was significantly decodable, with earliest onset and highest peak, in the Visual network (Figure S2A). Auditory stimulus information was significantly decodable, with earliest onset and highest peak, in the Auditory network (Figure S2B). These content-appropriate network results further highlight the high spatial precision achieved through our rigorous source modeling pipeline.

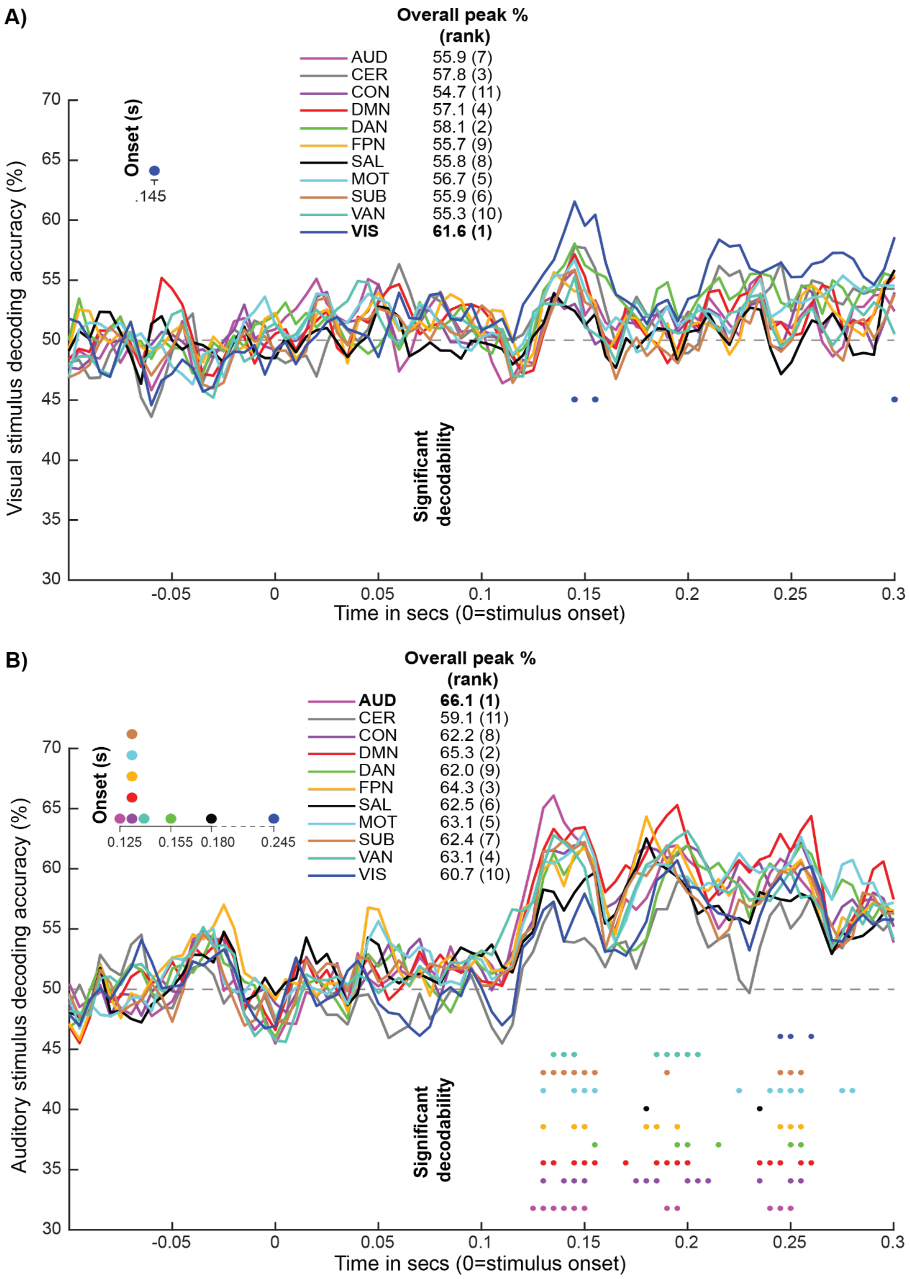


**Figure S2. Decoding stimulus categories within each sensory modality yields appropriate network peaks. A)** Decoding of visual stimulus categories (horizontal vs vertical) and **B)** auditory stimulus categories (low vs high pitch) across networks. Plotting conventions are the same as Figure 5 in the main manuscript.

**3. Stimulus-locked decoding clarifies the sequential timing of sensory and response information peaks.**

Figure S3A plots the network decoding of response information (left versus right hand) for the stimulus-locked trial data. Whereas the analyses in the main manuscript focused on response-locked data, this variation more easily illustrates the timing of sensory relative to response information peaks. As further validation of our source modeling precision, response information again peaked overall highest in the content-appropriate Motor network. The two-peak morphology seen in the response-locked data (Figure 5A) was again visible. To more easily visualize the relative timing, Figure S3B plots the sensory and response information timecourses for their respective peak networks (Visual and Motor) after zscore normalization, with the average reaction time also plotted. This highlights a sequential processing cascade from the early sensory decoding peak (S1) at 0.175s, to the later sensory peak (S2) at 0.265s, to the subsequent early motor peak (M1) which onset during sensory decoding and peaked at 0.340s, and finally the later motor peak (M2) which onset just prior to response commission and peaked at 0.570s. The gradual rise in response information during early sensory processing, culminating in M1's response peak just after the later sensory peak, is consistent with an evidence accumulation process that transforms sensory into response information to prepare a motor response. This is followed by execution of that response at M2 (and possibly feedback), which onset just prior to the trial reaction time. The timing of the stimulus-locked decoding peaks therefore supports linking of the 2-peak response decoding morphology to early motor preparation and late motor execution/feedback processes respectively.

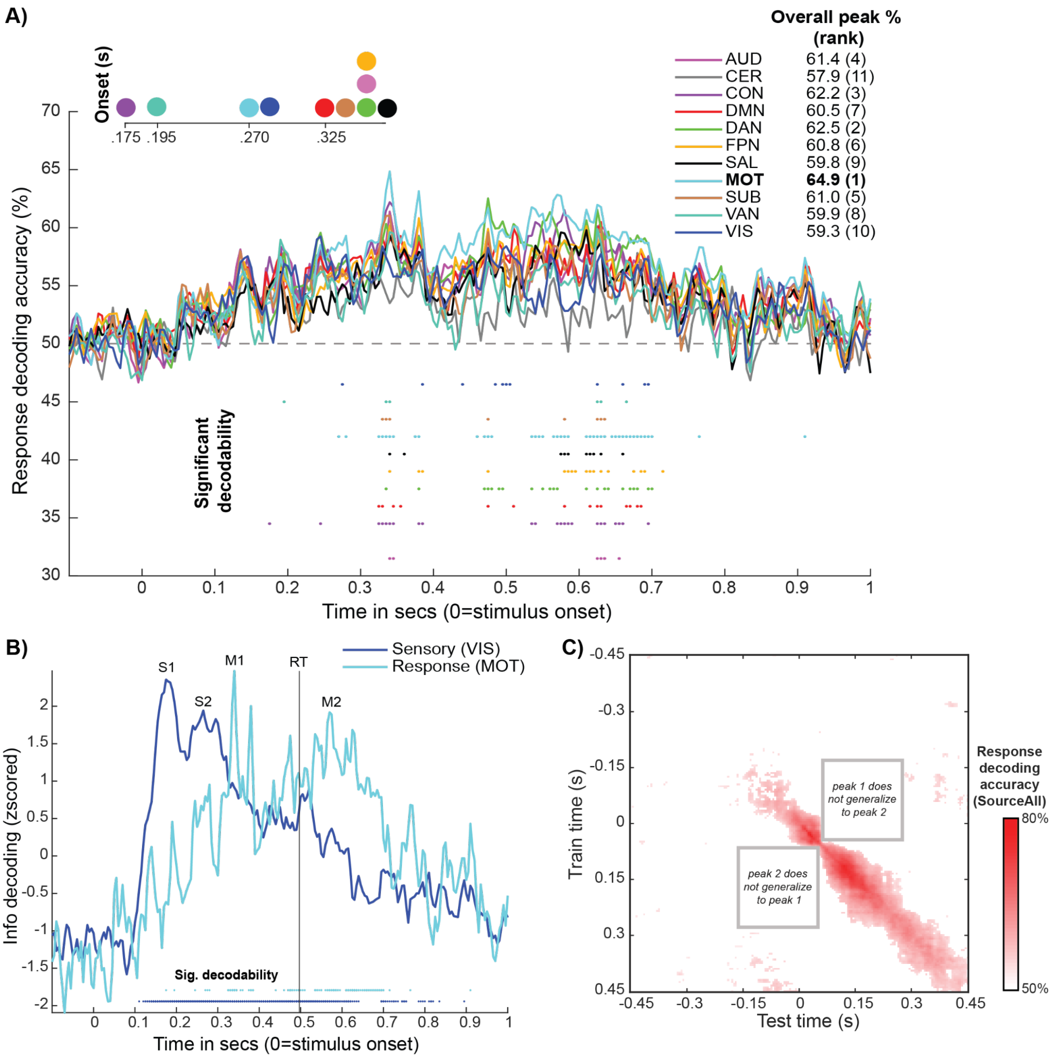


**Figure S3. Stimulus-locked decoding and temporal generalization analyses support transition from early motor preparation to later motor execution/feedback. A)** Stimulus-locked decoding of response information across networks. Plotting conventions are the same as Figure 5 in the main manuscript. **B)** Sensory decoding in the peak Visual network overlaid with response decoding in the peak Motor network for the stimulus-locked data. Timecourses are zscored to aid visualization. Average reaction time (RT) is also depicted. **C)** Plotted is the group matrix for temporal generalization conducted for response decoding in the SourceAll data (response-locked), thresholded by Wilcoxon signrank tests conducted for each element against 50% chance ($p<.05$, FDR-corrected). Gray boxes highlight the lack of generalization for the multivariate representations underlying the peak 1 and peak 2 clusters, consistent with sequential engagement of distinct response representations.

**4. Temporal generalization results suggest sequential engagement of distinct response codes.**

Both source and sensor group timecourses in Figure 4 exhibited two decoding peaks, which might reflect distinct multivariate representations (neural codes) underlying separate motor preparation and motor execution/feedback processes described previously (Ames et al., 2019; Cheyne et al., 2006; Elsayed et al., 2016; Shibasaki and Hallett, 2006; Wascher and Wauschkuhn, 1996; Wolpert and Ghahramani, 2000). To assess whether these two decoding peaks were underpinned by two distinct multivariate representations, we performed a temporal generalization analysis in the SourceAll data (King and Dehaene, 2014). Extending the 10-fold cross-validation scheme used for the main dynamic decoding analyses, we trained linear SVM classifiers at each timepoint and tested them on all other trial timepoints. Averaging over all folds (10) and subtrial loops (10) generated a training timepoint (181) x testing timepoint (181) matrix of classification accuracies for each subject. These matrices were averaged over subjects and thresholded by contrasting the element-wise classification accuracies against 50% chance via Wilcoxon signrank tests (with FDR-correction for multiple timepoint x timepoint comparisons).

Inspection of this matrix (Figure S3C) revealed two distinct clusters of response decodability corresponding to peak 1 (onsetting prior to response commission at ~-0.150 and generalizing to 0.075s) and peak 2 (onsetting at ~0.075s and generalizing late into the trial) in the response decoding timecourse. Critically, the lack of significant decodability on the off-diagonal of each peak cluster (gray boxes in Figure S3C) confirmed that each was underpinned by distinct multivariate representations. The timing of the representations is consistent with sequential engagement of motor preparation and motor execution processes described in animals, with the latter potentially also involving post-movement feedback (Ames et al., 2019; Elsayed et al., 2016; Wolpert and Ghahramani, 2000).

**5. Confining the network decoding to unique signals in each network does not change the pattern of results.**

As an alternative to the network decoding analyses in the main manuscript (Figure 5), we explored an approach that sought to remove common sources of signal variation across networks prior to performing dynamic MVPA. The motivation was to confine the decoding of response information to more "unique" network signals, and see if this affected the spatiotemporal pattern of results. For a given held-out region, we regressed out the 1st principal component estimated separately for regions within each network, excluding that held-out region and its affiliated network. Critically, the within-network regions were first demeaned to ensure that the 1st principal component captured each network's activation pattern rather than the network mean, as the former is more likely to drive the MVPA classifiers. Multiple linear regression models then fit the component scores for the 10 held-out networks (X) to each region's trial activation timeseries (Y), with the residuals then submitted to network decoding as before.

Figure S4 depicts the resulting unique network information timecourses. Whilst these appear more dissimilar than those in the main manuscript, we nevertheless find significantly decodable information in 10 out of the 11 networks (the Cerebellar network was not significantly decodable). This was observed despite the principal component regression successfully reducing variance associated with the common network signals, as indexed by the Pearson correlation between each region's activation timeseries before and after the correction (r=~.30 across regions). The average pairwise Pearson correlation between the group information timecourses (capturing unscaled variance explained) decreased from r=.94 to r=.69, and the average coefficient of determination (capturing scaled variance explained) also decreased from $R^2$=61.2% to $R^2$=8.71%. The fact that the majority of networks still contained significantly decodable response information despite these large reductions in shared variance strengthens the inference that task information is encoded in distributed fashion in the brain (Kauvar et al., 2020; Siegel et al., 2015).

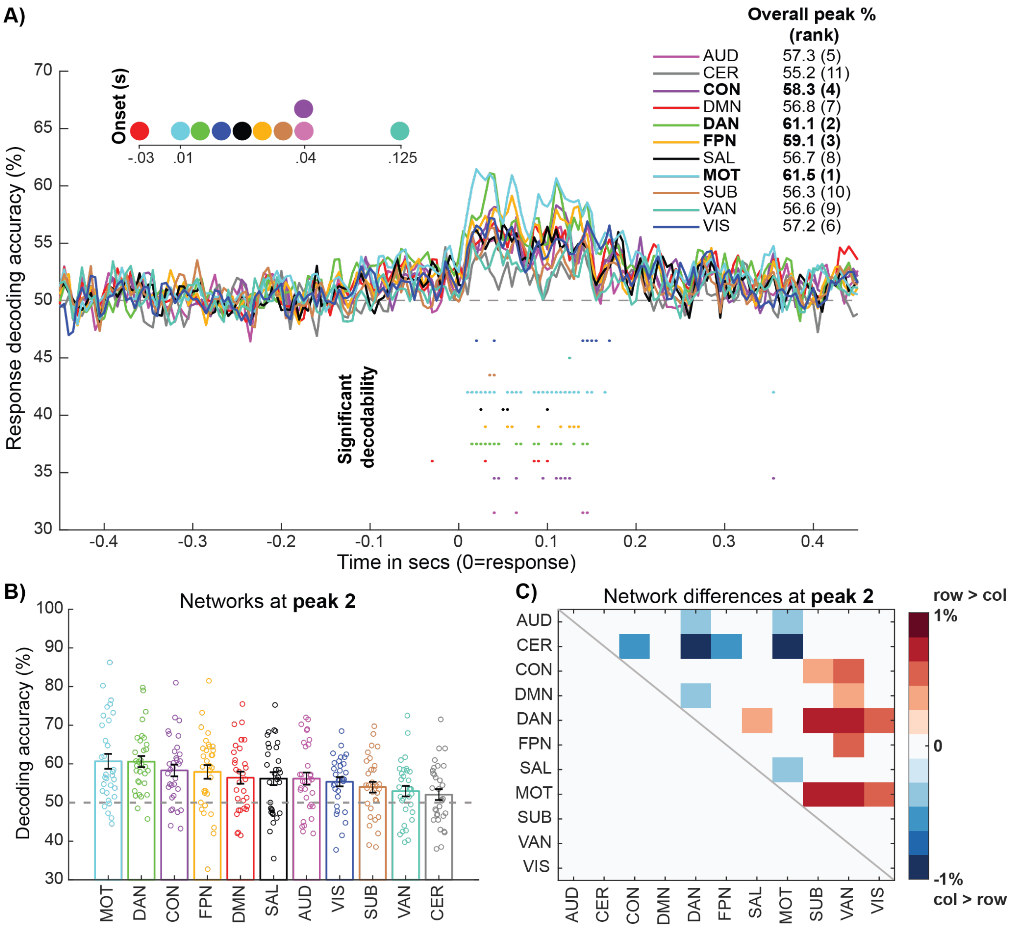


**Figure S4. Decoding of response information from unique network activation patterns. A)** Group decoding timecourses for the analysis that confined MVPA to unique network signals, color-coded by network affiliation. **B)** Response decoding accuracy ranked across networks at peak 2 (0.11s). **C)** Matrix capturing the significance of cross-network differences in decoding accuracy at peak 2 (via Wilcoxon tests, p<.05 FDR-corrected). Plotting conventions are the same as Figure 5 in the main manuscript.

The graded pattern of response information peaks across networks was also preserved. The content-appropriate Motor network again peaked highest overall, with the CCNs ranked next-highest. The 2-peak timecourse morphology was also largely preserved, as was the dynamic network coding profile of the CCNs. At the early peak 1 (0.02s), the Motor, DAN and Visual networks had the highest decodability. This pattern shifted at the later peak 2 (0.11s), wherein the Motor network and the 3 CCNs represented response information most strongly. Critically, as in the main analyses, decodability in the CCNs at peak 2 was significantly higher than most of the other functional networks (Figure S4C), but did not differ amongst themselves even at a relaxed threshold (p<.05 uncorrected).

The fact that the content-appropriate Motor network specialization and the early/late dynamic CCN segregation held in this unique network decoding analysis suggests these results are robust to analytic variation. Even though the pattern of results was well preserved, the absolute peak decodability for all networks was reduced e.g. the Motor network reduced from a peak of 79.8% in the main analyses to 61.4%. To avoid collateral effects of this overall diminished decoding response, we applied our main dynamic activity flow modeling analyses to the regular (non-unique) task timeseries.

**6. Analysis of conventional Pearson correlation restFC shows recovery of canonical functional network architecture in source EEG.**

Figure S5A plots the group averaged matrix for restFC estimated via conventional Pearson correlation. Inspection of the group matrix reveals visible on-diagonal network clustering typical of the canonical functional network architecture. For comparison, Figure S5B plots a group averaged fMRI restFC matrix, estimated from the "100 unrelated subjects" subset of the publicly released Human Connectome Project (HCP) Young Adult dataset (Van Essen et al., 2013). The similarity of the two group matrices was quantified via Pearson correlation computed on the upper diagonals, revealing a significant positive correlation (r=.17, p<.0001 via non-parametric Mantel test). Previous reports that directly compared source EEG/MEG and fMRI restFC matrices are lacking, but the present magnitude of similarity is comparable to that of a recent report (r=~.15; Wirsich et al., 2021). Significant similarity with the group fMRI matrix was maintained at the subject-level also, via Pearson correlation of the subject EEG and group fMRI matrices (mean r=.07, t(31)=10.3, p<.00001).

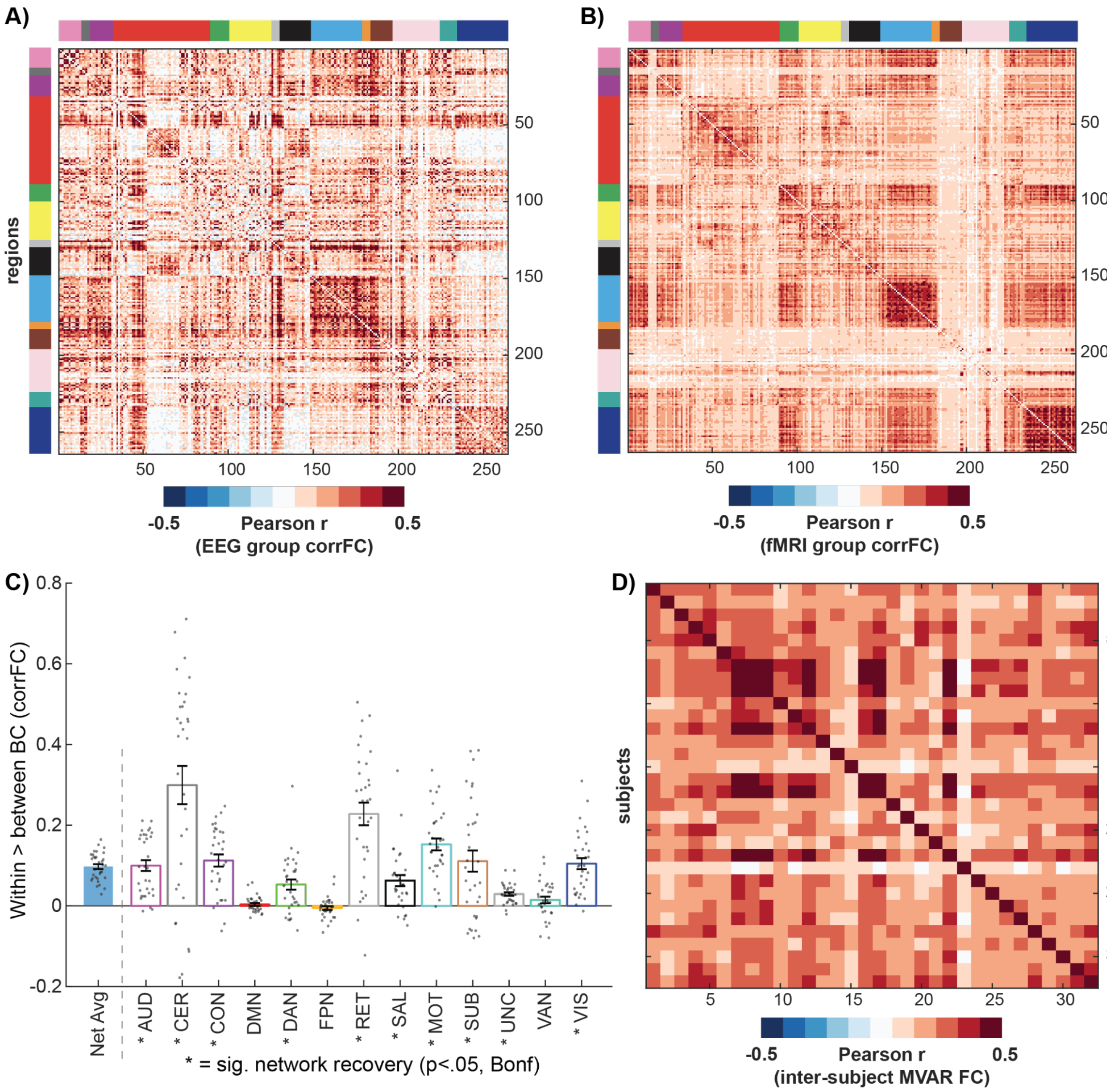


**Figure S5. RestFC estimation metrics (corrFC network recovery and MVAR FC reliability)**. **A)** Group averaged region-by-region matrix for field-standard Pearson correlation FC (corrFC) estimated in the present source EEG dataset. Colored panels link regions to their Power atlas network affiliations. Visible on-diagonal clustering suggests broadly good network recovery. **B)** Corresponding group matrix for fMRI corrFC estimated in the HCP dataset (n=100, see text for details). **C)** Graph theoretic assessment of network recovery (within-network minus between-network brain connectivity) in the source EEG data. The first “Net Avg” bar reflects the cross-network average of the graph measure, estimated in individual subjects (p<.00001). Asterisks denote significantly positive network recovery for individual networks (via one-sample ttest against 0 at p<.05, Bonferroni-corrected for multiple network comparisons). **D)** Inter-subject reliability of the MVAR restFC weights, computed as the Pearson correlation between all MVAR FC matrix elements pairwise across subjects. Mean inter-subject reliability r=.22 (standard deviation=.10, range=.01-.60).

To provide a more objective assessment of Pearson correlation-based network recovery, we computed a variation of the “global brain connectivity” (GBC; Cole et al., 2010) graph theoretic metric, which is also sometimes referred to as “unthresholded weighted degree centrality” (Rubinov and Sporns, 2011). This analysis formalized the assumptions underlying visual assessment of network recovery based on the degree to which on-diagonal (within-network) clustering in the FC matrix is greater than the off-diagonal (between-network). We hence averaged within-network and between-network (i.e. out-of-network) brain connectivity separately for each of the 13 Power atlas networks. Subtracting within- minus between-network brain connectivity for each network and subject, and contrasting these values against 0 via one-sample ttest, hence quantified the extent to which each functional network was reliably recovered in the FC matrix. The results of this graph theoretic analysis are presented in Figure S5C, revealing broadly positive network recovery in these source EEG data (i.e. network recovery was significant in the cross-network average, and for 10 out of the 13 individual networks at p<.05, Bonferroni-corrected for multiple network comparisons).

Overall, these results highlight that the standard correlation-based restFC architecture typically seen in fMRI data was broadly recovered in our source EEG data. Note that the dynamic activity flow modeling method that is the main focus of this paper uses a different and more advanced (from a causal connectivity standpoint) MVAR approach to restFC estimation (see Methods for details on these advanced features, and see Figure S5D for the inter-subject reliability of MVAR restFC). Hence, our intention in providing these correlation-based FC results is to serve as a general frame of reference to guide the still nascent field of source EEG/MEG functional connectivity. To this end, the reported success of dynamic activity flow modeling provides a compelling demonstration of the utility of our MVAR restFC approach, given that this network modeling framework goes beyond the above more conventional and primarily descriptive (Mill et al., 2017) FC analyses. This is achieved by embedding the FC weights in a generative network modeling framework that is applied empirically to simulate the emergence of cognitive task information underlying behavior. Whereas we extend the modeling approach to test alternative network models of behavior in the present report (see lesioning analyses in Figure 7), recent work involving standard (non-dynamic) activity flow modeling has adapted the approach to adjudicate between alternative methods of FC estimation (McCormick et al., 2022). Hence while it is reassuring that certain canonical features of correlation-based FC are recovered in our data (e.g. greater within- versus between-network brain connectivity), we find the reported success of dynamic activity flow modeling in predicting task behavior even more compelling.

**7. Full analyses with non-causal temporal filtering.**

Figure S6 provides the full set of results after applying non-causal temporal filtering during preprocessing, rather than the causal filtering used for the main results. These results were included to provide some basis for comparing the timing of the reported information/activation effects with previous reports that are more likely to have applied non-causal filtering. However, as we note in the Method, the dynamic activity flow modeling results after non-causal filtering are presented with the caveat of potential temporal leakage from target to-be-predicted Motor regions (at t0) to the lagged predictor terms (t0-n lags). This potential was entirely avoided by use of causal filtering in the main analyses (Figure 6), albeit at the cost of introducing a delay in the filtered output. Whilst the non-causal filter accounts for this delay, we still recommend caution when interpreting the absolute timing of these results, given the potential for such filters to overcorrect for the delay (shifting onsets of signals prior to when they actually occur; de Cheveigné and Nelken, 2019).

Figure S6 demonstrates that the overall pattern of results held after non-causal filtering (see Figure S6 legend for a summary). However, some differences in temporal morphology were apparent. Firstly, the onset latencies for the dynamic decoding and the motor ERP activation effects were earlier for the non-causal compared to causal filtering results (see Figure S6 caption for key onset differences). This is anticipated by the backward pass of the non-causal filter that attempts to correct for the onset delay introduced by the forward pass (the causal filter only applies a forward pass; de Cheveigné and Nelken, 2019; Widmann and Schröger, 2012). However, we also observed a less anticipated change in the shape of the dynamic decoding and motor ERP timecourses: whereas the main causal filter results yielded a consistent 2-peak shape, the non-causal filter results yielded a single peak centered around response commission. Evidence from the temporal generalization extension of the dynamic MVPA analysis (Figure S3C) and the statistically reliable variation in network engagement across the two response decoding peaks (see descriptive analyses in Figure 5 and more explanatory “lesioning” analyses in Figure 7) suggested that the 2-peak morphology obtained for the causal filter reflected separate motor preparation and motor execution multivariate representations. The presence of these two subcomponents underlying response representation has also been reliably established in animal electrophysiological studies (Ames et al., 2019; Elsayed et al., 2016). The possibility therefore presents that the non-causal filter effectively blurred two representationally distinct decoding peaks into one. This might have arisen from the backwards pass of the non-causal filter pushing low frequency signals backwards and forwards in time, which risks introducing distortions in output EEG waveforms as previously highlighted (see Chapter 12 in Luck, 2005). The potential for such distortion further substantiates our treatment of the non-causal filter results as supplementary to the causal filter results reported in the main manuscript. Critically, it is worth emphasizing that the high accuracy of dynamic activity flow modeling held across both analyzed filter types.

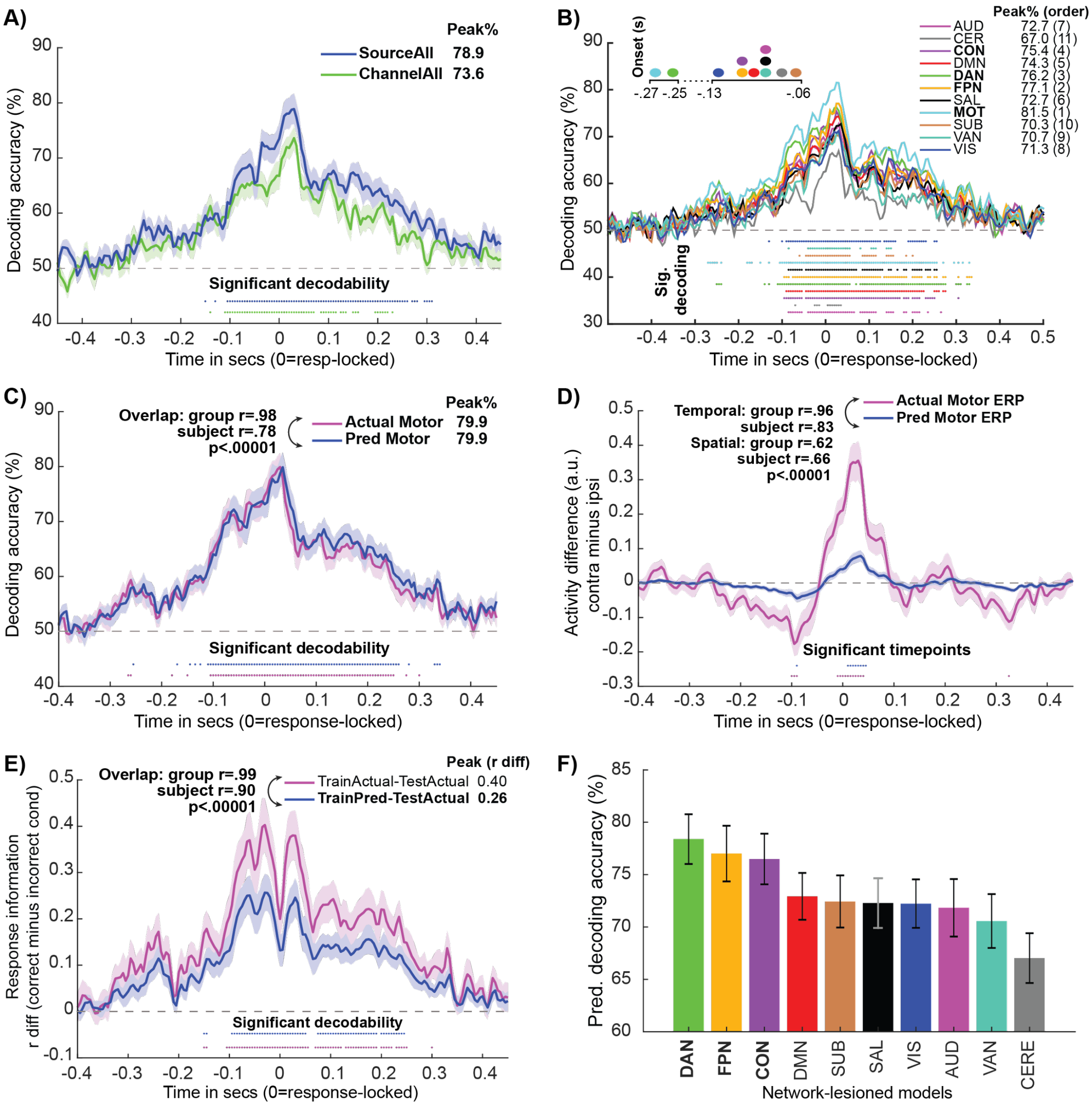


**Figure S6**. **Full pattern of results holds after applying non-causal temporal filtering during preprocessing. A)** Using all source-modeled regions as features (SourceAll) yields higher overall response decodability than using all EEG sensors as features (SensorAll). Plotting and statistical conventions are the same as Figure 4. SourceAll onset=0.105s earlier for non-causal compared to causal. **B)** Network decoding reveals prominent roles of the Motor network and the CCNs in representing response information. Same conventions as Figure 5A. Motor network onset=0.135s earlier for non-causal. **C)** Dynamic activity flow modeling yields highly accurate predictions of response information dynamics in the Motor network. Same conventions as Figure 6A. Predicted decoding onset=0.155s earlier for non-causal. **D)** The model accurately captures the underlying motor ERP activation effects (same conventions as Figure 6B; predicted timecourse onset=0.080s earlier for non-causal), and **E)** the fine-grained representational geometry (same conventions as Figure 6C; TrainPred-TestActual timecourse onset=0.060s earlier for non-causal). **F)** Network lesioning variant of dynamic activity flow modeling highlights strong causal influence of CCNs on response information flow to the Motor network (extracted at cross-network group time-to-peak at 0.030s; same conventions as Figure 7B/D).

### 8. Effect of field spread regression in improving dynamic activity flow model performance.

As detailed in the Methods section, we took a number of steps to rigorously rule out influences of field spread artifacts on the dynamic activity flow modeling results. These included use of beamformer source modeling, distal spacing of source regions localized from the Power atlas (Power et al., 2011) and exclusion of contemporaneous (t0) source predictor terms from the dynamic activity flow modeling step. We also regressed out each to-be-predicted Motor target's task timeseries from all source predictors prior to running the main dynamic activity flow modeling analyses. For these supplementary analyses, we re-ran the modeling after excluding this regression step, so as to quantify its impact on model accuracy. Figure S7 shows that inclusion of this step actually numerically improved model prediction accuracy across an exhaustive set of metrics. We theorize that this resulted from the regression step removing artifactual field spread influences (known to be contemporaneous/instantaneous; Schoffelen and Gross, 2009; Stinstra and Peters, 1998) shared by the targets and predictors at each timepoint, thereby uncovering the lagged influences that veridically underpin task information flow. To demonstrate, consider the activity of a predictor region i1 at timepoint t0-1 ($i1_{t0-1}$); the activation state at this timepoint is likely influenced both by artifactual field spread shared with a to-be-predicted region j1 at the same timepoint ($j1_{t0-1} \rightarrow i1_{t0-1}$), as well as by its own lagged communication to target j1 at future timepoint t0 ($i1_{t0-1} \rightarrow j1_{t0}$). The field spread regression acted to remove the former artifactual contemporaneous influence ($j1_{t0-1} \rightarrow i1_{t0-1}$), leaving greater scope to pick up the latter's contribution to lagged communication comprising real neural activity flow ($i1_{t0-1} \rightarrow j1_{t0}$). These findings hence highlight important considerations for network modeling of source EEG/MEG data.

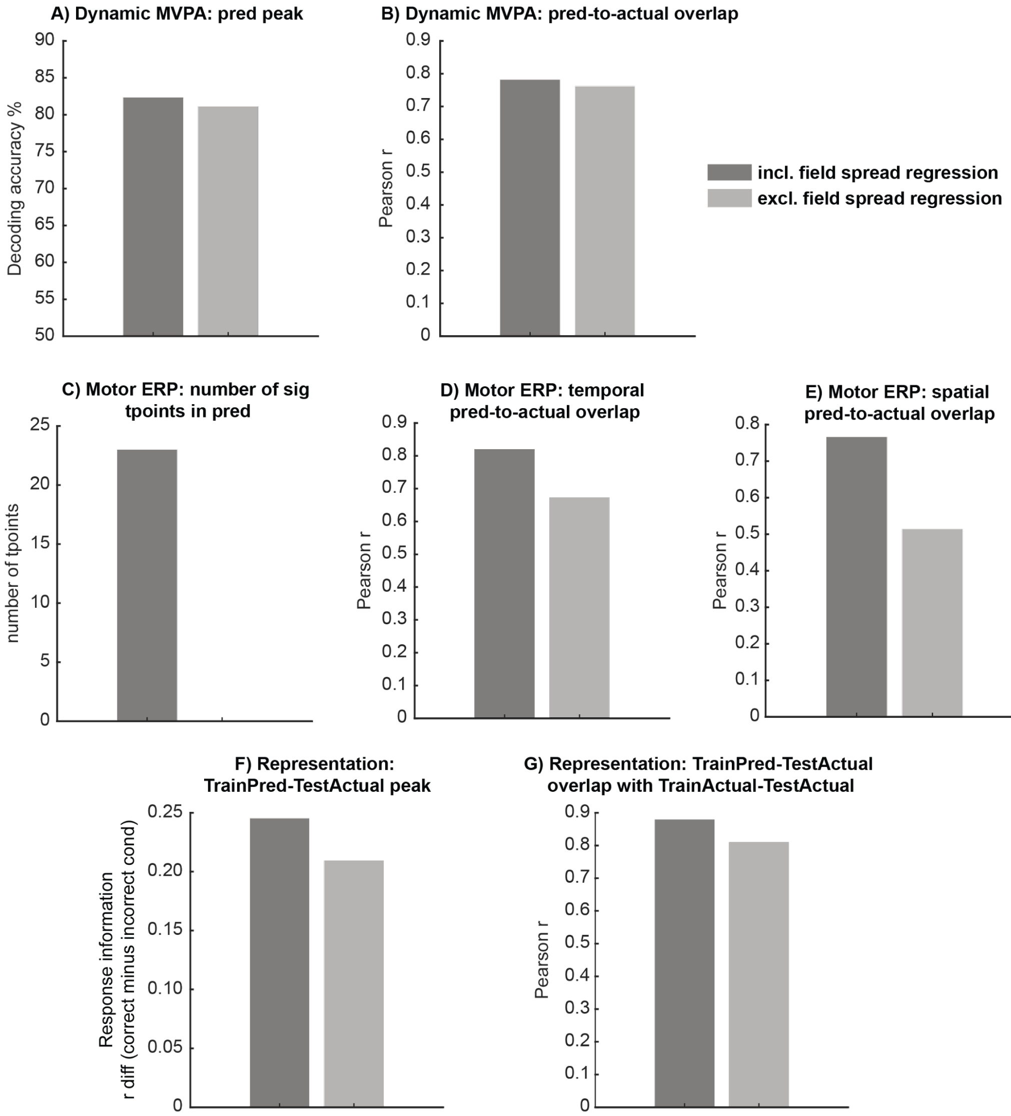


**Figure S7. Improved dynamic activity flow model performance with inclusion of field spread regression step.** Bar plots depict comparisons of model performance for inclusion (dark gray) versus exclusion (light gray) of this step across multiple metrics: A) dynamic MVPA predicted response decoding peak, B) dynamic MVPA predicted-to-actual timecourse overlap (subject-level), C) motor ERP number of significant timepoints in the predicted difference wave, D) motor ERP predicted-to-actual timecourse temporal overlap (subject-level), E) motor ERP predicted-to-actual regional activation pattern spatial overlap (subject-level), F) representational overlap predicted decoding peak (TrainPred-TestActual), G) representational overlap predicted-to-actual timecourse overlap (TrainPred-TestActual with TrainActual-TestActual).

### 9. Alternative metric for network lesioning results: accuracy in predicting subject-level response decoding timecourses.

As an alternative means to probe the generative contributions of the functional networks to response information flow in the simulated lesioning analysis, we also contrasted the subject-level response decoding prediction accuracy (i.e. Pearson r overlap between the model-predicted and actual response timecourses) across the "lesioned" network models. The results in Figure S8A again highlight strong contributions from the CCNs in predicting individualized response information dynamics in the Motor network, with these networks having the top 3 prediction accuracies overall.

The CCN contributions also significantly differed from each other, with the DAN yielding significantly higher prediction accuracy than the FON and CON (FDR-corrected p<.05, Figure S8B), whereas the CON and FPN did not significantly differ even at a lenient p<.05 uncorrected threshold. This recapitulates the strong generative influence of the CCNs with an alternative model performance metric to the predicted decoding accuracy (%) measure used in the main manuscript.

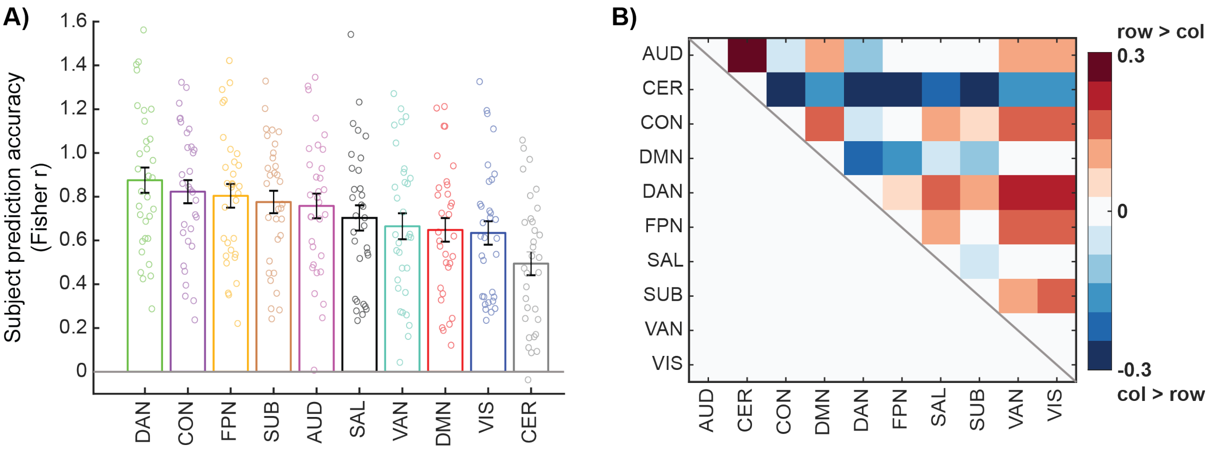


**Figure S8. Comparison of predicted-to-actual response decoding timecourse overlap for network-lesioned models again highlights prominent CCN influence. A)** Overlap (Fisher-z transformed Pearson r) ranked across network models, revealing highest prediction accuracy for the CCN network models. Each bar represents the mean and standard error for each network, with overlaid subject data points. **B)** Matrix capturing cross-network differences in decoding timecourse overlap. Plotted is the pairwise difference in subject overlap (Fisher-z transformed r values), thresholded via paired Wilcoxon tests (p<.05, FDR-corrected across pairwise network comparisons). Positive values denote significantly higher overlap for the row network > the column network, and vice versa for the negative values.
